# The noncoding circular RNA *circHomer1* regulates synaptic development and experience-dependent plasticity in mouse visual cortex

**DOI:** 10.1101/2024.07.19.603416

**Authors:** Kyle R. Jenks, Ying Cai, Marvin Eduarte Nayan, Katya Tsimring, Keji Li, José C. Zepeda, Gregg R. Heller, Chloe Delepine, Jennifer Shih, Shiyang Yuan, Yao Zhu, Ye Wang, Yangyang Duan, Amy K. Y. Fu, Taeyun Ku, Dae Hee Yun, Kwanghun Chung, Nikolaos Mellios, Mriganka Sur, Jacque Pak Kan Ip

## Abstract

Circular RNAs (circRNAs) are a class of closed-loop, single stranded RNAs whose expression is particularly enriched in the brain. Despite this enrichment and evidence that the expression of circRNAs are altered by synaptic development and in response to synaptic plasticity *in vitro*, the regulation by and function of the majority of circRNAs in experience-dependent plasticity *in vivo* remain unexplored. Here, we employed transcriptome-wide analysis comparing differential expression of both mRNAs and circRNAs in juvenile mouse primary visual cortex (V1) following monocular deprivation (MD), a model of experience-dependent developmental plasticity. Among the differentially expressed mRNAs and circRNAs following 3-day MD, the circular and the activity-dependent mRNA forms of the *Homer1* gene, *circHomer1* and *Homer1a* respectively, were of interest as their expression changed in opposite directions: *circHomer1* expression increased while the expression of *Homer1a* decreased following 3-day MD. Knockdown of *circHomer1* delayed the depression of closed-eye responses normally observed after 3-day MD. *circHomer1*-knockdown also led to a reduction in average dendritic spine size prior to MD but critically there was no further reduction after 3-day MD, consistent with impaired structural plasticity. *circHomer1*-knockdown also prevented the reduction of surface AMPA receptors after 3-day MD. Synapse-localized puncta of the AMPA receptor endocytic protein Arc increased in volume after MD but were smaller in *circHomer1*-knockdown neurons, suggesting that *circHomer1* knockdown impairs experience-dependent AMPA receptor endocytosis. Thus, the expression of multiple circRNAs are regulated by experience-dependent developmental plasticity, and our findings highlight the essential role of *circHomer1* in V1 synaptic development and experience-dependent plasticity.

**Significance Statement:** Circular RNAs (circRNAs) are a class of closed-loop RNAs formed through back-splicing of exon and/or intron junctions. Initially considered as byproducts of aberrant RNA splicing with limited function, recent studies have implicated circRNAs in various neurological disorders. Despite their abundant expression in the brain, the role of circRNAs in experience-dependent plasticity remains poorly understood. We conducted an *in vivo* transcriptome analysis of circRNAs whose expression was regulated by experience-dependent plasticity in the mouse visual cortex and highlight *circHomer1*, a circRNA derived from the *Homer1* gene, as a circRNA critical for synaptic development and experience-dependent plasticity *in vivo*.

## Introduction

Experience-dependent plasticity is a tightly regulated process that requires orchestrated gene transcription to form mRNAs that are subsequently translocated and translated into proteins to induce the functional and structural reorganization of synapses(1). Though we now know a great deal about the mRNAs recruited by and required for experience-dependent plasticity, it remains unclear if other forms of non-coding/coding RNAs are as tightly regulated as their mRNA counterparts and if they contribute to experience-dependent plasticity(2, 3). Recent RNA-sequencing screens have shown that a plethora of genes that produce alternatively spliced mRNAs also generate close-looped isoforms composed of backspliced and covalently-joined exons and/or introns known as circular RNAs (circRNAs)(4). Although much is still unknown about the precise mechanisms that govern the biogenesis, regulation, and function of circRNAs; there is emerging evidence that circRNAs may have important roles in the brain(3, 5). They are dynamically expressed during brain development, are localized to synapses(6), and a small number of circRNAs have been shown to affect neuronal gene expression, miRNA availability, and regulate behavior(7–9). However, it remains unclear how many circRNAs might play a role in regulating experience-dependent plasticity, highlighting the need for screens to identify putative candidates.

Ocular dominance plasticity in mouse binocular primary visual cortex (V1) is a well-defined model of cortical plasticity induced by monocular deprivation (MD), typically via suture of the eye-lids, during a developmental critical period(10–15). During this window, which in mice begins at postnatal day (P)21 and ends around P35(16), MD induces a reduction of V1 responses to the deprived eye after ∼3 days of deprivation, followed by an increase of responses to the non-deprived eye(17) after ∼7 days of deprivation(11, 13–15, 18–20). MD also leads to the activity-dependent up- and down-regulation of many mRNAs known to regulate synaptic plasticity(10–15). We have successfully used MD in the past to screen for and identify miRNAs, another unique class of RNA species, regulated by and critical for experience-dependent plasticity(2). Thus, we sought to similarly use MD to identify circRNAs in mouse V1 that could be involved in regulating experience-dependent plasticity and synaptic function.

By screening for differentially expressed mRNAs and circRNAs in the developing mouse V1 following 3 days of MD (3-day MD), we discovered multiple circRNAs whose expression was regulated by experience-dependent plasticity. We further explored the role of the circRNA *circHomer1* in experience-dependent plasticity, as the expression of *circHomer1* was opposite that of its linear, activity-dependent form *Homer1a* and *circHomer1* has previously been shown to be localized to synapses and involved in cognitive flexibility and reversal learning(9, 21, 22). Using *circHomer1*-specific knockdown in V1, we showed that *circHomer1* delays normal closed-eye response depression following 3-day MD. While *circHomer1* knockdown alone decreases the average size of dendritic spines on layer 2/3 neurons, it also prevented the decrease in spine size following 3-day MD. Knockdown of *circHomer1* also impaired downregulation of the AMPA receptor subunit GluA1 following 3-day MD, putatively by reducing Arc protein expression at dendritic spines. Together, these data provide the first evidence of an experience-dependent circRNA that is required for normal synaptic development and experience-dependent plasticity induced by short-term MD.

## Results

### circRNA expression is regulated by experience-dependent plasticity

To identify potential experience-regulated circRNAs, we performed MD in mice beginning at P25, collected tissue from V1 and examined the transcriptomic profile after 3-day MD (Fig. 1A). We normalized RNA expression in the hemisphere contralateral to the deprived eye, where the majority of change in synaptic drive occurs, to expression in the ipsilateral hemisphere (within-subject) which receives significantly less input from the deprived eye and has significantly less change in gene expression(2). Before exploring the circRNA expression profile, we first examined the mRNA expression profile to validate our results against previously observed mRNA regulation by MD. We identified a total of 1173 differentially-expressed mRNAs, with more mRNAs downregulated than upregulated (472 upregulated and 701 downregulated, Fig. 1B). mRNAs known to be downregulated during MD, such as *Bdnf* and *Nptx2*(23), were also downregulated in our data (Supplementary Table 1). Having confirmed that our preparation replicated the known regulation of mRNA expression by 3-day MD(24), we next examined the regulation of circRNAs. We used a published circRNA alignment tool, circtools, to detect circRNA transcripts by virtue of their unique out-of-order junctional reads(23). A total of 1489 unique circRNAs were detected, with 73 differentially-expressed (Fig. 1C, Supplementary Table 2). Of the differentially-expressed circRNAs, we observed 27 significantly upregulated circRNAs and 46 downregulated circRNAs. Some of the genes from which these differentially-expressed circRNAs were derived have known roles in synaptic plasticity and function (Supplementary Fig. 1A). Indeed, we also identified changes in the corresponding mRNA isoforms of several differentially expressed circRNAs. Of these, the linear isoforms of *circHomer1*, *circPrkce*, and *circDcun1d4* were of interest as they exhibited changes in the opposite direction compared to their circular counterparts, suggesting their circRNA expression was regulated by experience independently of the overall gene transcription rate (Fig. 1D). We also performed a transcriptomic analysis of mRNAs and circRNAs in V1 after 7-day MD (Supplementary Fig. 1B-E and Supplementary Tables 3 and 4). *circPrkce*, *circDcun1d4*, and their linear isoforms were not differentially expressed at this timepoint, while linear *Homer1a* remained downregulated and *circHomer1* was no longer upregulated. Based on these findings, and the known role of *Homer1a* in synaptic plasticity, we chose to more closely examine the experience-dependent and developmental regulation and role of *circHomer1* in mouse V1.

**Figure 1.**
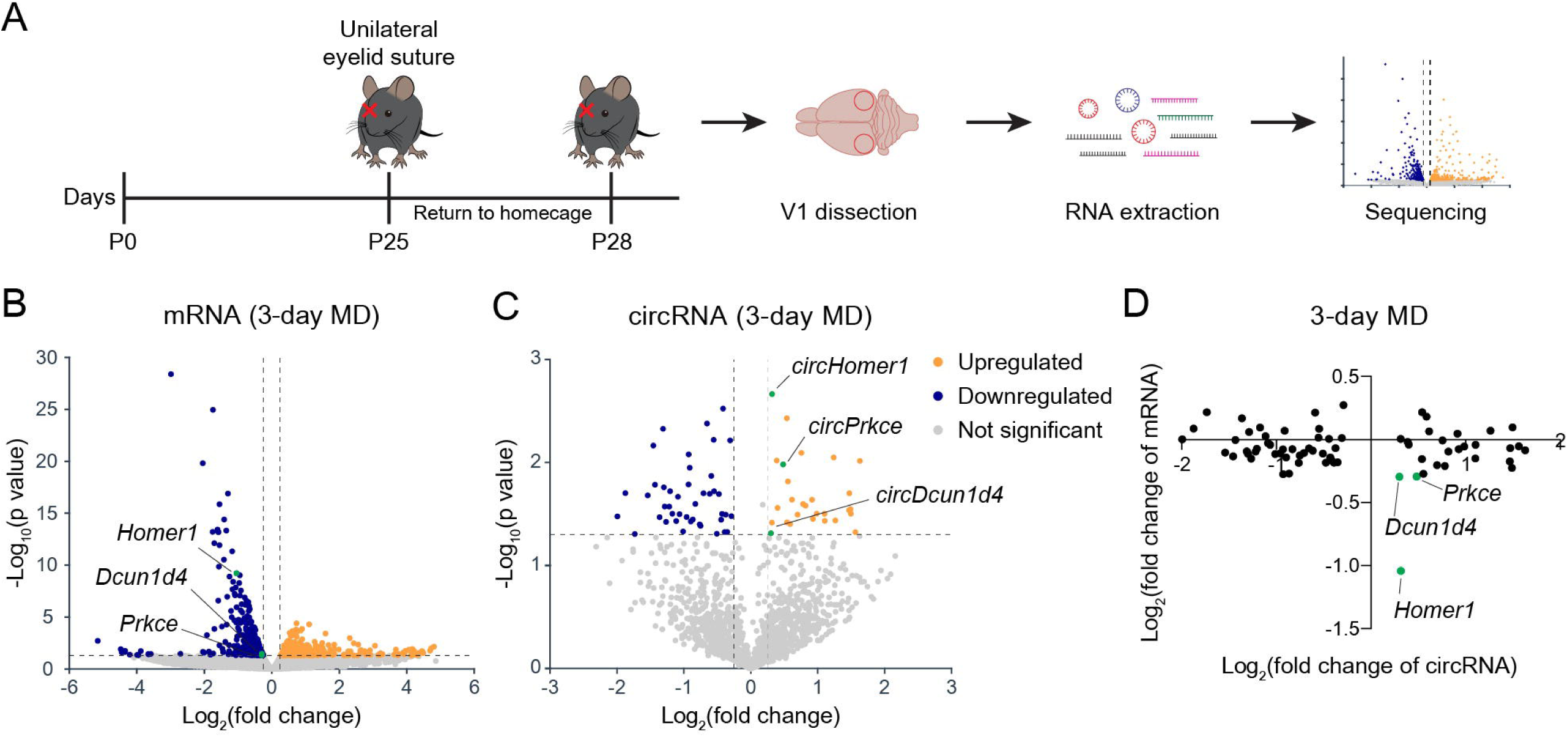
Experience-dependent circRNAs in V1 identified through MD. (A) Timeline and schematic of 3-day MD RNAseq screen. (B) Volcano plot of mRNA sequencing of V1 after 3-day MD (n = 3 mice per group). (C) Volcano plot of circRNA sequencing of V1 after 3-day MD (n = 3 mice per group). The ipsilateral hemisphere (ipsi) served as a control for MD induced changes in the contralateral hemisphere (contra) for both mRNA and circRNA expression. (D) Fold change of differentially-expressed circRNAs and their mRNA isoforms after 3-day MD. The mouse and brain silhouette were adapted from https://doi.org/10.5281/zenodo.8044766 and https://doi.org/10.5281/zenodo.3925971.

### *circHomer1* expression is developmentally regulated

*circHomer1* is composed of exons 2 through 5 of the *Homer1* gene, with backsplicing between exon 2 and 5 (Fig. 2A). Using primers targeting the backsplice junction of *circHomer1*, we next sought to validate the observed changes in contralateral V1 *circHomer1* expression, normalized to ipsilateral V1, after 3-day and 7d-day MD using RT-qPCR (Fig. 2B). In addition, we examined several mRNAs known to be regulated by MD. As expected after 3-day MD, *Bdnf* and *Nptx2* mRNA were downregulated in contralateral V1(24). *Homer1a* mRNA was also significantly downregulated after 3-day MD, in contrast to *circHomer1*, validating the dissociation between *Homer1a* and *circHomer1* expression after 3-day MD (Fig. 2B, left). After 7-day MD, *Stat1*(25), a regulator of homeostatic synaptic plasticity, was upregulated in contralateral V1 (Fig. 2B, right) as previously shown for prolonged MD(17, 24, 25). We also saw that the expression of *circHomer1* was significantly decreased after 7-day MD along with *Homer1a* (Fig. 2B, right). These results validate the RNAseq data showing that experience-dependent regulation of *circHomer1* expression is distinct at 3-day MD from that of its linear experience-dependent counterpart *Homer1a*.

**Figure 2.**
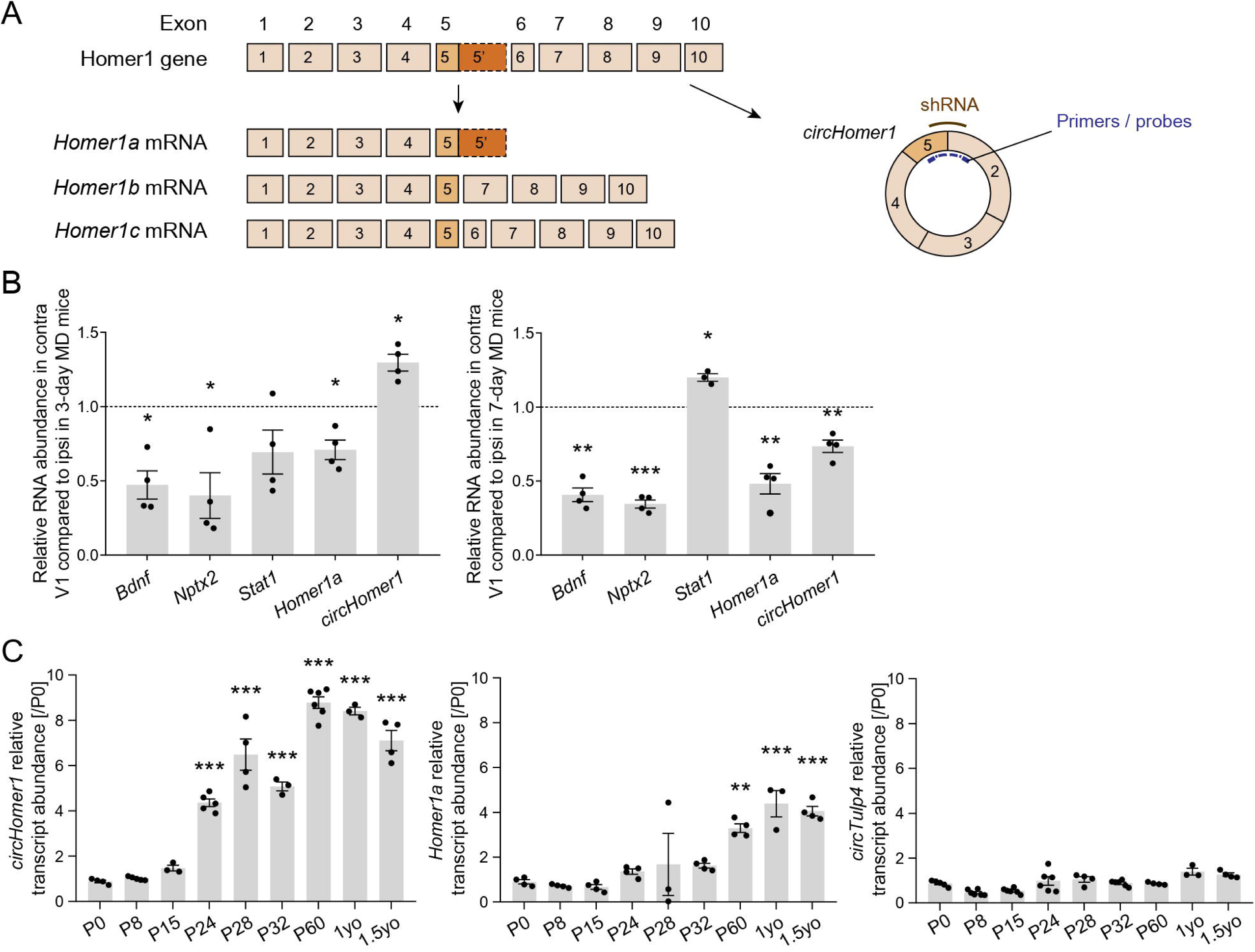
*circHomer1* expression is regulated by experience-dependent plasticity and upregulated during the ocular dominance critical period. (A) The mouse Homer1 gene encodes long and short versions of the synaptic protein Homer1 and a noncoding circRNA, *circHomer1*. (B) RT-qPCR quantification of selected mRNAs and *circHomer1* after 3-day or 7-day MD (n = 3-4 mice per group, paired t-test). (C) *circHomer1* (left), *Homer1a* (middle), and *circTulp4* (right) expression measured by RT-qPCR at several timepoints in mouse V1 across development and into adulthood (n = 3-6 mice per group, one-way ANOVA following Tukey multiple comparisons). Data are presented as mean ± SEM. *p < 0.05, **p < 0.01, ***p < 0.001.

Expression of many mRNAs required for normal, developmental experience-dependent plasticity peak during the critical period. To determine the developmental expression profiles of *Homer1a* and *circHomer1* over the mouse’s lifespan, we measured their expression in V1 at various time points using RT-qPCR. *circHomer1* expression increased significantly starting at ∼P24, aligning to the start of the critical period (Fig. 2C, left), while *Homer1a* expression did not increase significantly until P60, past the close of the critical period (Fig. 2C, middle). *circHomer1* expression appeared to plateau at P60 and remained at similar levels of expression in mice as old as 1.5 years. Expression of another circRNA, *circTulp4*, did not change across development or adulthood indicating that the developmental expression profile of *circHomer1* is not a shared feature of all circRNAs (Fig. 2C, right).

### *circHomer1*-depletion delays ocular dominance plasticity

The regulation of *circHomer1* expression by MD and the peak in *circHomer1* expression at the start of the critical period mirrors that of linear mRNAs with known roles in ocular dominance plasticity. To determine if *circHomer1* is required for ocular dominance plasticity following MD, we used a short hairpin RNA (shRNA)-mediated depletion strategy targeting the backsplice junction of *circHomer1*, which has been previously validated *in vivo* to specifically downregulate *circHomer1*, and not linear *Homer1* mRNAs (including *Homer1a*, *Homer1b*, and *Homer1c*)(9), in the cortex. We re-confirmed the specificity of knockdown using adeno-associated viruses (AAV)-mediated transduction of the shRNA in mouse V1, which resulted in a significant reduction of *circHomer1* RNA with no changes in *Homer1a*, *Homer1b*, or *Homer1c* mRNA expression (Supplementary Fig. 2A). sh-circHomer1 also had no effect on total Homer1 protein levels (Supplementary Fig. 2B).

To examine the effects of *circHomer1* depletion on ocular dominance plasticity, we quantified eye-specific responses in the binocular zone of V1 by using optical imaging of intrinsic hemodynamic signals measured before and after 3-day or 7-day MD (Fig. 3A). We then compared results between mice injected with a sh-scramble (control) or sh-circHomer1 virus (Fig. 3B). To compare the relative visual drive elicited by the two eyes in V1, the normalized difference between contralateral (dominant-eye) and ipsilateral responses during optical imaging was measured to determine the ocular dominance index (ODI). Change in ODI following closure of the contralateral eye (MD) was then used as a measure of experience-dependent plasticity(26).

**Figure 3.**
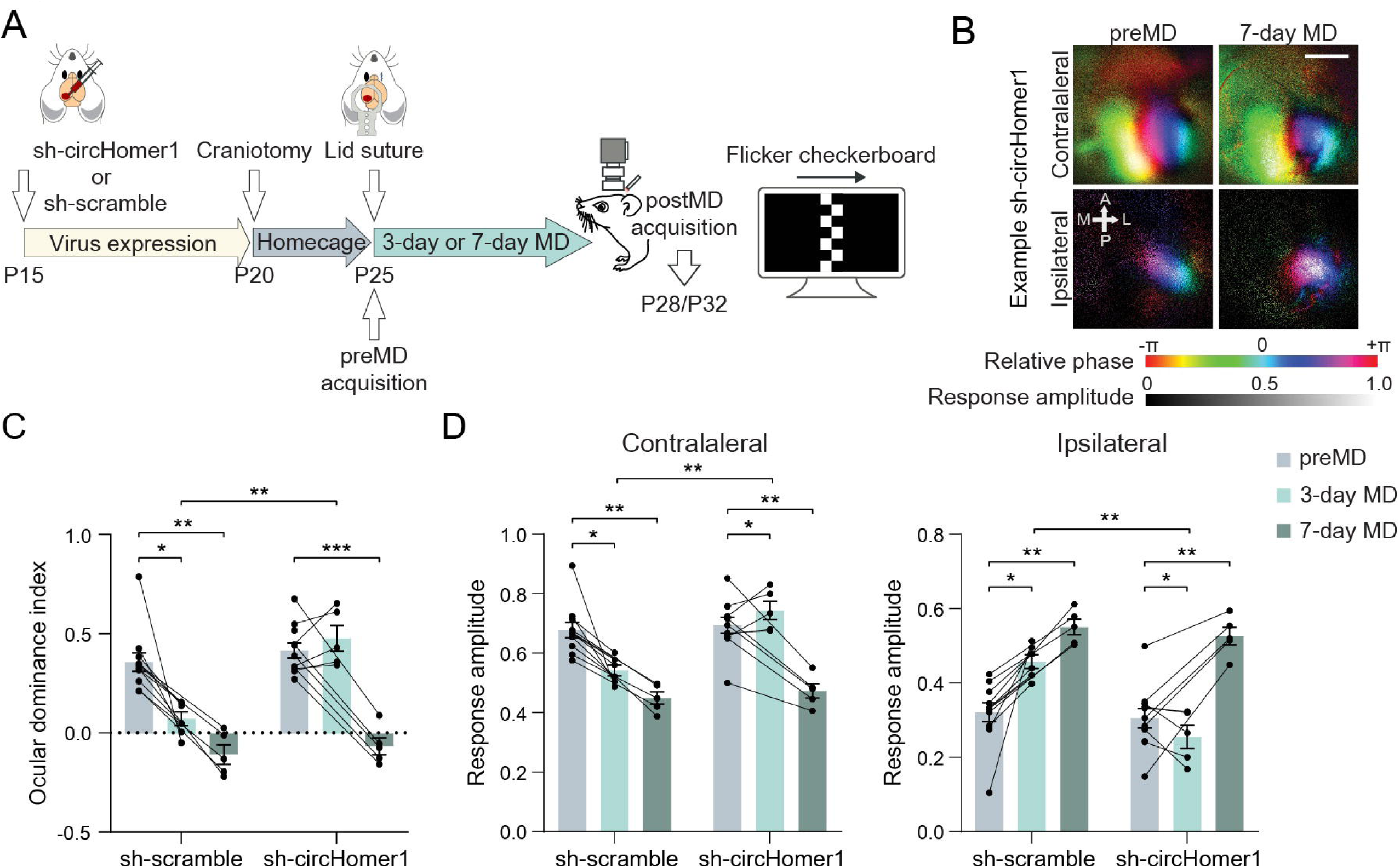
*circHomer1* depletion delays the expression of ocular dominance plasticity in V1 following MD. (A) Mice were injected with either sh-circHomer1 or sh-scramble virus at P15, a craniotomy was performed at P22 to implant a cranial window, then a pre-MD optical imaging session was done and the contralateral eye-lid sutured at P25. After either 3-day or 7-day MD, the eyelid was reopened and post-MD imaging done. (B) Example retinotopic maps from a sh-circHomer1 mouse, obtained with optical imaging. The color corresponds with different phases of the visual stimulus (consistent with the visual field map) and brightness shows the amplitude of cortical response. Top, contralateral closed-eye responses, pre-MD (left) and after 7-day MD (right). Bottom, ipsilateral open-eye responses. Scale bar = 500 µm. (C) Ocular dominance index (ODI) changes after 3-day or 7-day MD for mice injected with sh-scramble or sh-circHomer1 virus. Each dot shows the average ODI for one animal, and grey lines show the change of ODI for one animal (n = 5-11 mice per group, mixed-effects model following Holm-Sidak multiple comparisons test). (D) Normalized response amplitudes driven by the contralateral (left) and ipsilateral (right) eyes (n = 5-11 mice per group, mixed-effects model following Holm-Sidak multiple comparisons test). Data are presented as mean ± SEM. *p < 0.05; **p < 0.01; ***p < 0.001.

Prior to MD (preMD), there was no difference in baseline ODI between sh-scramble and sh-circHomer1 mice, with responses biased towards the dominant, contralateral eye (Fig. 3C). After 3-day MD, the ODI of sh-scramble mice decreased significantly (shifted towards the open, ipsilateral eye) as expected(24, 27, 28). In contrast, the ODI of sh-circHomer1 mice were unchanged after 3-day MD (Fig. 3C). After 7-day MD, however, the ODI of sh-circHomer1 mice were significantly decreased and no longer significantly different than that of sh-scramble mice (Fig. 3C).

The ODI shifts at 3-day and 7-day MD are driven by an initial decrease in contralateral drive and delayed increase in ipsilateral drive, respectively(24, 27, 28). To see if both these eye-specific changes instead occurred between 3-day and 7-day MD in sh-circHomer1 mice, we examined the responses driven by the contralateral and ipsilateral eyes separately. Unlike in the sh-scramble mice, and as expected from the lack of change in ODI, contralateral closed-eye responses of sh-circHomer1 mice did not decrease after 3-day MD. Similarly, ipsilateral open-eye responses were already significantly increased in sh-scramble mice after 3-day MD whereas they were paradoxically reduced in sh-circHomer1 mice (Fig. 3D). Despite the differences in eye-specific changes after 3-day MD, however, by 7-day MD both contralateral and ipsilateral responses in the sh-circHomer1 mice were comparable to the sh-scramble mice (Fig. 3D). Thus, *circHomer1* depletion does not grossly impair the eye-specific development of contralateral and ipsilateral visual drive and delays, but does not block, the expression of binocular experience dependent plasticity following MD.

### *circHomer1* regulates dendritic spine morphology

MD induces rapid remodeling of the dendritic spines of mouse layer 2/3 V1 neurons, with 3-day MD significantly decreasing average spine volumes(10). As we had found that depletion of *circHomer1* blocked the depression of closed eye responses at 3-day MD (Fig. 3), we hypothesized that depletion of *circHomer1* would likewise impair 3-day MD-induced spine shrinkage. We injected an AAV expressing RFP and either sh-circHomer1 or sh-scramble shRNA into the binocular region of V1 at P15, prior to the start of the critical period. We also injected AAV9-hSyn-DIO-EGFP (structural marker) and AAV9-CaMKIIa-Cre virus in order to sparsely label neurons and quantified spine morphology of labeled layer 2/3 V1 neurons from mice that had undergone no MD (No MD), 3-day MD, or 7-day MD. To achieve a more detailed analysis of the dendritic spine structure than is possible with traditional confocal imaging, we used epitope-preserving magnified analysis of the proteome (eMAP) to expand our tissue samples (Fig. 4A). The apical dendrites of V1 layer 2/3 pyramidal neurons expressing both GFP and RFP were selected for further analysis (Fig. 4B, left). Based on their morphology, dendritic spines were classified into four categories: mushroom, stubby, thin, and filopodia using standard criteria (Fig. 4B-C)(29).

**Figure 4.**
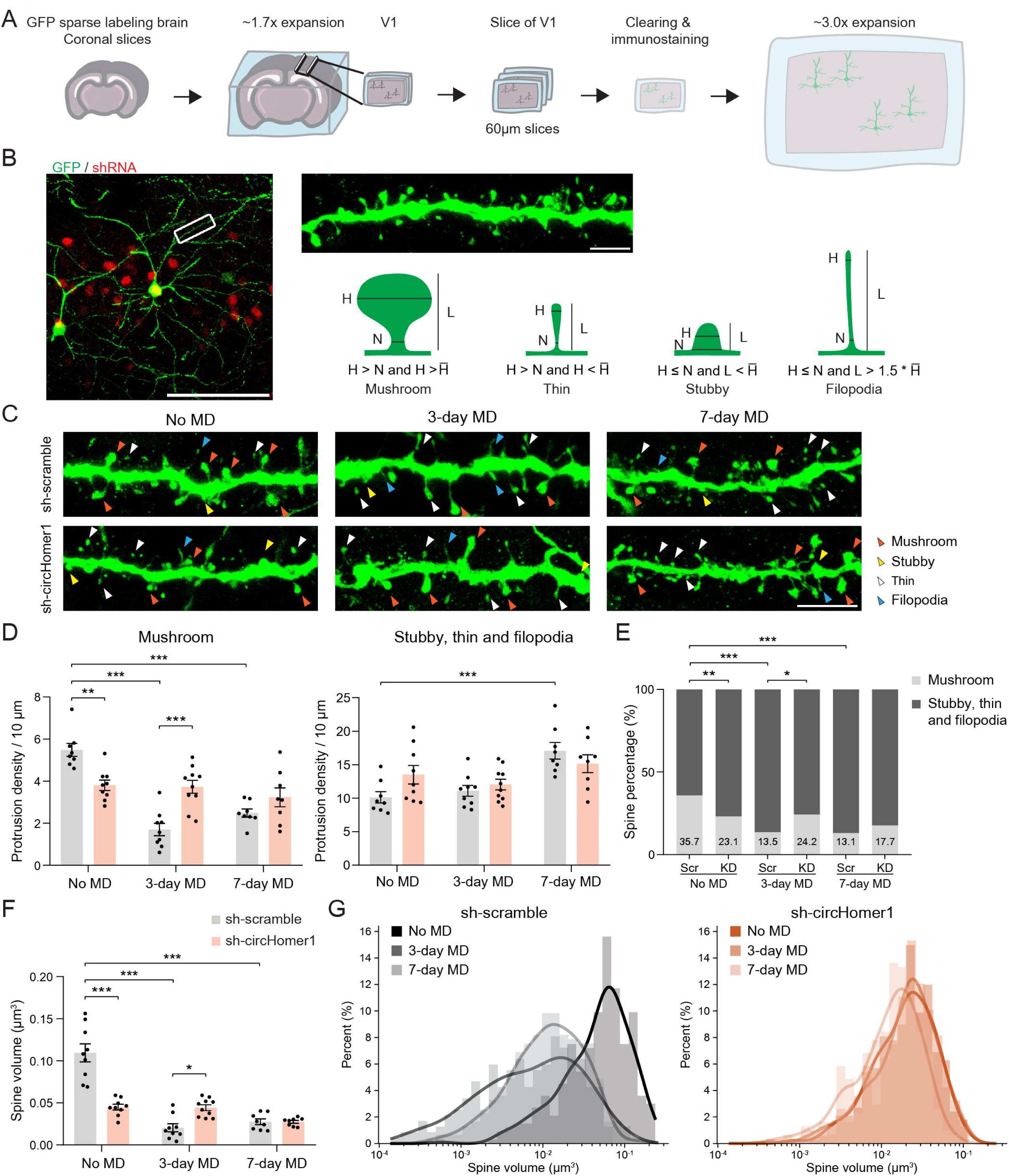
Depletion of *circHomer1* affects spine morphology of V1 neurons. (A) A schematic of tissue preparation and eMAP. (B) Neurons expressing both GFP and RFP (denoting shRNA) were selected for analysis (left). Dendritic spines were classified into four categories based on their morphology (right). Scale bars = 100 µm (left, estimated to be 33.33 µm prior to 3x expansion), and 10 µm (right, estimated to be 3.33 µm prior to 3x expansion). H, head width; N, neck width; L, length; H̅, average head width in No MD sh-scramble group. (C) Representative images showing apical dendrites of layer 2/3 neurons in V1 in the No MD, 3-day MD, and 7-day MD condition from the sh-scramble (top) or sh-circHomer1 (bottom) group. Arrows indicate different types of dendritic spines. Scale bar = 10 µm (estimated to be 3.33 µm prior to 3x expansion). (D) Density of mushroom spines, and immature spines (stubby spines, thin spines and filopodia) on apical dendrites of layer 2/3 neurons in V1 in the No MD, 3-day MD, and 7-day MD condition from the sh-scramble (grey) or sh-circHomer1 (pink) group (n = 8-10 dendrites from 3 mice per group, values were normalized to 3x expansion factor, two-way ANOVA following Tukey multiple comparisons). (E) Spine morphology types as a percentage of total spines. The percentage of mushroom spines are labeled on the bar. (F) Volume of dendritic spines on apical dendrites (same data as in D and E, values were normalized to 3^3^ expansion factor, two-way ANOVA following Tukey multiple comparisons). (G) Distribution of spine volume in sh-scramble and sh-circHomer1 mice in the 3 conditions (n = 235-334 dendritic spines from 3 mice per group). Data are presented as mean ± SEM. *p < 0.05; **p < 0.01; ***p < 0.001.

In No MD mice, sh-circHomer1 led to a significant drop in the density and percentage of mature mushroom spines(30), but not immature spines (stubby, thin, and filopodia)(30) (Fig. 4D-E). This basal effect was surprising given the normal baseline ODI and eye-specific responses we had observed in preMD sh-circHomer1 mice. After 3-day MD in sh-scramble mice, as expected, there was a significant decrease in the density and percentage of mature mushroom spines when compared to the No MD condition. Conversely, the density of both mature and immature spines remained unchanged after 3-day MD in the sh-circHomer1 mice. After 7-day MD, in the sh-scramble mice the density and percentage of mature spines remained lower compared to the No MD condition, while the density of immature spines showed a significant increase. Mature and immature spine density in sh-circHomer1 mice after 7-day MD again remained unchanged compared to the No MD condition, but was no longer significantly different than the sh-scramble mice (Fig. 4D-E).

As categorical classification could obscure more subtle changes in the dendritic spines of sh-circHomer1 neurons, we also analyzed the average spine volume by performing 3D reconstruction. *circHomer1* depletion led to a more than 50% decrease in average spine volume in the No MD condition (Fig. 4F, Supplementary Fig. 3A). After 3-day MD, the average spine volume decreased by more than 70% in sh-scramble mice compared to the No MD condition. In contrast, 3-day MD did not significantly alter spine volume in sh-circHomer1 mice (Fig. 4F-G, Supplementary Fig. 3B). After 7-day MD, spine volume was still significantly decreased in sh-scramble mice compared to the No MD condition, and while the sh-circHomer1 spine volume remained unchanged, the sh-scramble and sh-circHomer1 spine volumes were not significantly different at 7-day MD (Fig. 4F, Supplementary Fig. 3C). Taken together, our findings suggest that *circHomer1* plays a crucial role in regulating the structural morphology of dendritic spines both during normal development and following short-term MD.

Considering the effects *circHomer1* depletion had on the morphology of synapses in the No MD condition, we sought to determine how *circHomer1* depletion affected the electrophysiological properties of neurons. We used whole-cell patch clamp recordings in acute slices from P28-33 mice to measure passive membrane and synaptic properties of layer 2/3 neurons infected with either sh-circHomer1 or sh-scramble virus. We did not find any difference in baseline neuronal excitability (Supplementary Fig. 4) or the amplitude of miniature excitatory postsynaptic currents (mEPSC; Supplementary Fig. 5C), but there was a decrease in mEPSC frequency (Supplementary Fig. 5D), consistent with a decrease in protrusion density(31). While the basal structural and electrophysiological deficits following *circHomer1* depletion were surprising given the apparently normal binocular visual drive observed prior to MD, experience-dependent plasticity following MD was delayed as opposed to simply occluded by *circHomer1* depletion. We therefore sought to further investigate the mechanism by which *circHomer1* depletion impaired experience-dependent plasticity.

### *circHomer1* controls surface AMPA receptor expression

The major mechanism underlying both reduced closed-eye drive and spine shrinkage following MD is a reduction in surface AMPA receptors(25–27). Therefore, we investigated whether *circHomer1* depletion disrupted surface AMPA receptor trafficking following MD using a biotinylation assay of the AMPA receptor subunit, GluA1 (Fig. 5A). As expected, 3-day MD led to a significant reduction of surface GluA1 compared to No MD in sh-scramble mice. There was no reduction of GluA1 levels after 3-day MD in sh-circHomer1 mice (Fig 5A). By 7-day MD, however, there was no significant difference in surface GluA1 expression between sh-scramble and sh-circHomer1 mice. Based on these results, we hypothesized that *circHomer1* was necessary for normal activity-dependent AMPA receptor endocytosis. To examine this further, we treated primary neurons *in vitro* with bicuculline, a GABA_A_R antagonist, to induce activity-dependent AMPA receptor internalization(32). 1- or 2-days of treatment with bicuculline induced upregulation of *circHomer1* in neurons (Supplementary Fig. 6A), and *circHomer1*-depletion impaired bicuculline induced endocytosis of surface AMPA receptors visualized via SEP-GluA1 expression (Supplementary Fig. 6B,C). These data demonstrate that *circHomer1* depletion impairs experience- and activity-dependent changes in surface GluA1 expression levels, thus indicating a potential functional mechanism for the disrupted ocular dominance plasticity after 3-day MD.

**Figure 5.**
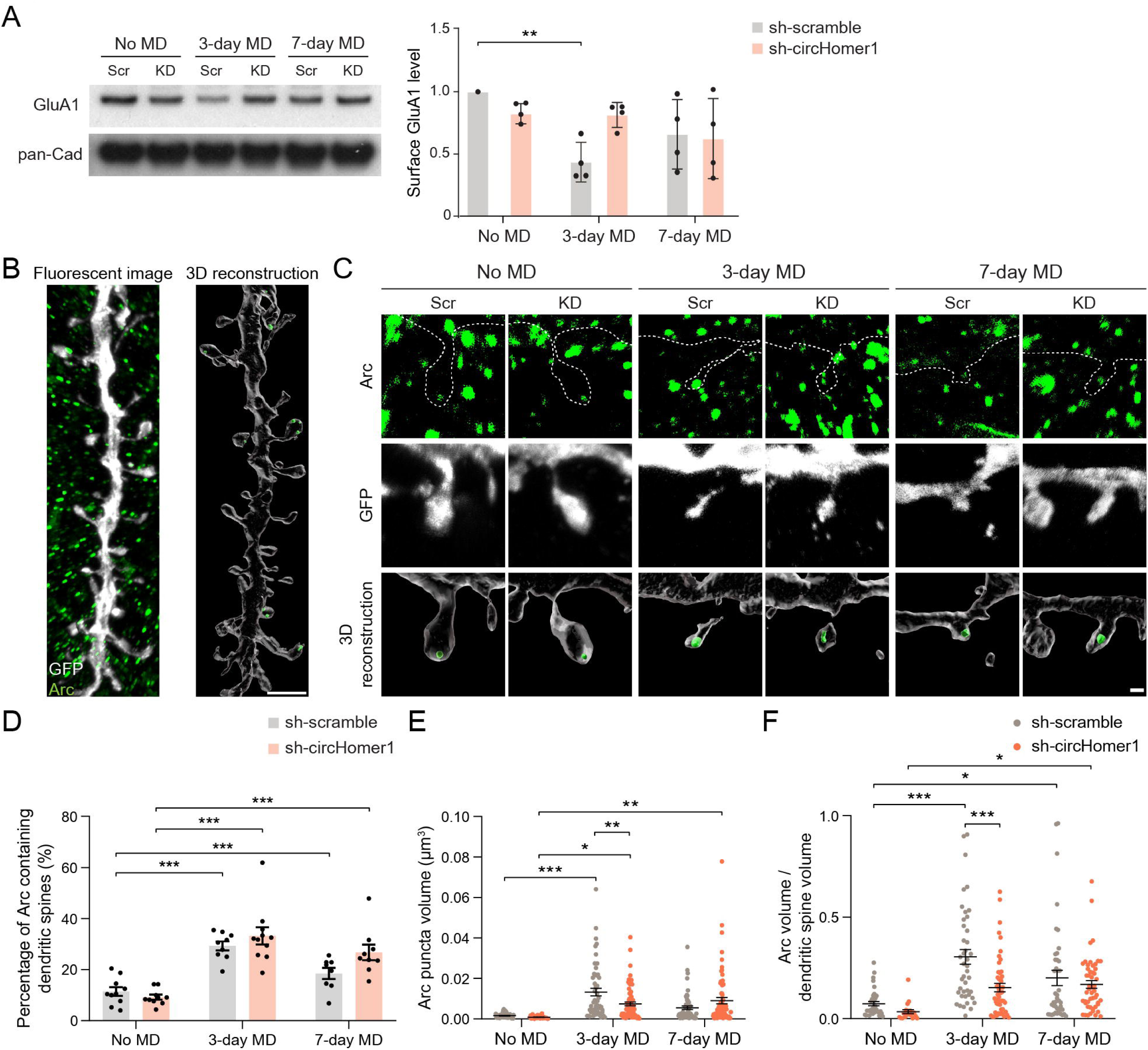
Trafficking of surface AMPA receptors following MD is regulated by *circHomer1*. (A) Left: Western blot of cell surface expression of GluA1 in V1 of mice infected with lentivirus after MD. Scr, sh-scramble; KD, sh-circHomer1. Right: Quantification of surface GluA1 level (n = 4 mice per group, two-way ANOVA following Tukey multiple comparisons). (B) Representative images showing 3D reconstruction of a segment of dendrites staining with GFP and Arc after eMAP processing. Scale bar = 5 µm (estimated to be 1.67 µm prior to 3x expansion). (C) Representative images of 3D-reconstructed Arc puncta at dendritic spines of V1 neurons after MD. Scale bar = 1 µm (estimated to be 0.33 µm prior to 3x expansion). (D) Percentage of dendritic spines containing Arc puncta (n = 8-11 dendrites from 3 mice per group, two-way ANOVA following Tukey multiple comparisons). (E) Arc puncta volume at dendritic spines (n = 29-71 puncta from 8-11 dendrites from 3 mice per group, values were normalized to 3x expansion factor, two-way ANOVA following Tukey multiple comparisons). (F) The ratio of Arc puncta volume and corresponding dendritic spine volume (n = 20-51 dendritic spines from 8-11 dendrites from 3 mice per group, two-way ANOVA following Tukey multiple comparisons). Data are presented as mean ± SEM. *p < 0.05; **p < 0.01; ***p < 0.001.

How might a circRNA contribute to AMPA receptor endocytosis? One possibility, given the previous observation of dendritically localized *circHomer1*(33) and binding to linear *Homer1b* mRNA(21), is through regulating other dendritically localized mRNAs critical to AMPA receptor endocytosis(22). The activity-regulated cytoskeleton-associated protein (Arc) is an immediate early gene product that has been shown to play a critical role in AMPA receptor endocytosis(32, 34), and *Arc* mRNA is trafficked to dendrites and translated locally in response to synaptic activity. We therefore tested whether Arc protein expression at dendritic spines is affected by *circHomer1* depletion following MD. We again injected AAV expressing RFP and either sh-scramble or sh-circHomer1, and sparsely labeled neurons in V1 with GFP as described in Figure 4. After processing the tissue with eMAP, postsynaptic Arc protein was labeled by immunostaining, and pyramidal neurons in layer 2/3 of V1 expressing both RFP and GFP were selected for analysis. Next, we performed 3D reconstruction of the imaged dendrites (Fig. 5B), and quantified the volume of Arc puncta expressed at dendritic spines (Fig. 5C). Intriguingly, the percentage of Arc-containing spines and the volume of Arc puncta increased significantly after 3- and 7-day MD in both sh-scramble and sh-circHomer1 mice as compared to the No MD conditions (Fig. 5D,E). However, the mean volume of Arc puncta and the Arc volume/spine volume ratio in sh-circHomer1 neurons was significantly lower than that in sh-scramble neurons after 3-day MD (Fig. 5E,F). There was no difference in the mean volume of Arc puncta and the Arc volume/spine volume ratio between sh-scramble and sh-circHomer1 neurons at 7-day MD. This is consistent with the impairment of surface GluA1 removal and spine shrinkage after 3-day MD in sh-circHomer1 mice as compared to control (Figs. 4, 5A). Together, these results demonstrate that *circHomer1*-depletion impairs GluA1 internalization following 3-day MD, which may be explained by *circHomer1*-depletion reducing the quantity of Arc protein accumulating at dendritic spines in response to MD.

## Discussion

circRNAs are abundantly expressed in mammalian brains, are preferentially derived from genes encoding synaptic proteins(33, 35), and several circRNA have recently described roles in synaptic plasticity or function(7–9, 21, 22). Based on these findings, we performed an unbiased screen to identify candidate, novel experience-dependent plasticity regulated circRNAs using *in vivo* linear- and circular-transcriptome analyses. We identified several promising circRNAs with differential expression in contralateral binocular V1 following 3-day MD in critical period age mice. We focused on *circHomer1*, a well-known circRNA derived from the *Homer1* gene which encodes important synaptic scaffolding proteins and an activity dependent linear isoform, *Homer1a*, which is a key regulator of glutamatergic synaptic signaling(36).

*circHomer1* and *Homer1a* were differentially regulated by 3-day MD, but in opposing directions with the expression level of *circHomer1* increasing and that of *Homer1a* decreasing following short-term MD. Following longer term, 7-day MD, however, expression levels of both *circHomer1* and *Homer1a* were significantly decreased (as measured via RT-qPCR). The developmental expression of *circHomer1* increased rapidly in V1 between P15 and P28, wherein the critical period for ocular dominance plasticity begins, which in combination with its differential expression following 3-day MD suggested a role for *circHomer1* in critical period plasticity. Indeed, loss of *circHomer1* disrupted ocular dominance plasticity following 3-day MD without effecting baseline visual responses. However, depletion of *circHomer1* alone also led to less mature dendritic spines and decreased mEPSC frequency. The normal, albeit delayed expression of ocular dominance plasticity following 7-day MD, however, suggests that the 3-day MD phenotype is not simply a result of occlusion. Indeed, while *circHomer1* depletion impaired 3-day MD induced spine shrinkage, AMPA receptor endocytosis, and accumulation of synaptic Arc protein these effects were not apparent when compared to their respective controls at 7-day MD.

The various proteins derived from the *Homer1* gene play critical roles in regulating synapse development, synaptic strength, and homeostatic synaptic scaling(37–41). At the postsynaptic density, the activity-dependent short protein isoform of the *Homer1* gene, Homer1a, has dominant-negative effects on the constitutive longer forms of Homer1 (Homer1b/c), and through this interaction regulates the clustering of mGluRs(38–40) and mGluR-dependent NMDA and AMPA receptor currents(38, 42–45). *circHomer1* appears to have a similar dominant negative effect as its linear, activity-dependent counterpart by instead binding directly to dendritic *Homer1b* mRNA which is necessary in the orbitofrontal cortex for normal reversal learning(21). Despite the fact that Homer1a and *circHomer1* both share activity dependent regulation and antagonize Homer1b expression at the synapse, their regulation by and role in developmental experience-dependent V1 plasticity are clearly distinct. In contrast to the upregulation of *circHomer1*, *Homer1a* is reduced following 3-day MD (Fig. 2C), and depletion of Homer1a does not affect the shift in ODI and reduction of contralateral, closed-eye responses as a result of 3-day MD(46). Homer1a depletion instead alters the establishment of contralateral bias of V1 responses under basal conditions, which is not affected by *circHomer1* depletion (Fig. 3C,D). Thus, although both Homer1a and *circHomer1* are regulators of cortical plasticity, they serve different roles in V1 development and experience-dependent plasticity.

It remains mechanistically unclear how neuronal activity regulates *circHomer1* expression, as the only demonstrated precursor of *circHomer1* is non-activity dependent *Homer1b* pre-mRNA(47). Given that activity-dependent *Homer1a* pre-mRNA contains the relevant regions to form *circHomer1*, including a section of intron 5(47), it is plausible that *circHomer1* could derive from *Homer1a* pre-mRNA, thereby allowing its expression to change in tandem with *Homer1a* mRNA even assuming a fixed ratio of linear/back-splicing, and thus circularization rate. However, during MD and development *in vivo* and bicuculline treatment *in vitro*, we observed timepoints where changes in *circHomer1* expression appear decoupled from or even opposing changes in *Homer1a*. Circular RNA species decay more slowly than their linear counterparts(48) and changes in transcription thus tend to result in larger, longer relative increases in circRNA levels(49), but this alone seems insufficient to explain this discrepancy. More plausible, perhaps, is that the circularization rate is itself regulated by activity(47).

Thus, we asked whether a constant circularization rate of *Homer1b* and *Homer1a* pre-mRNA, a variable circularization rate of *Homer1b* pre-mRNA, or a variable circularization rate of both *Homer1b* and *Homer1a* pre-mRNA best fit our observed data (Supplementary Fig. 7). Our models shows that across multiple paradigms, a constant circularization rate of *Homer1a* and *1b* pre-mRNA or a variable circularization rate of *Homer1b* pre-mRNA alone is insufficient to explain the observed expression levels of *circHomer1*. Instead, our data are best fit when adding a variable, activity-dependent circularization rate of both *Homer1b* and *Homer1a* pre-mRNA. In this way, *circHomer1* levels can increase when *Homer1a* mRNA levels do not change or even decrease. Importantly, the model accounts for the bidirectional change in *circHomer1* after 3-day versus 7-day MD. (Fig. 2C-D, Supplementary Fig. 7A). Such regulation could be facilitated by recently described molecular pathways underlying *circHomer1* biogenesis(47) or by changes in other *Homer1* mRNA products not measured in our experiments (for instance, *Homer1b*(21)). Future work is needed to determine if *Homer1a* pre-mRNA can indeed give rise to *circHomer1*, whether circularization of pre-mRNA occurs at a sufficient frequency to meaningfully decreases linear mRNA availability, and if circular RNA decay rate is static or itself regulated by activity as is the case for some mRNA species(7, 8, 50, 51).

An important conclusion of our study is that *circHomer1* has a critical role in the reduction of synaptic drive that accompanies the initiation of MD. Specifically, we found that *circHomer1* knockdown alters the time course of surface GluA1 trafficking and Arc protein expression during experience-dependent plasticity. However, the exact mechanism of *circHomer1* in synaptic development and plasticity warrants further investigation. Previous studies suggest that *circHomer1* regulates alternative splicing of mRNA isoforms(9) as well as synaptic expression of *Homer1b* mRNA in the orbitofrontal cortex via direction competition for HuD binding(21). Overall expression of several postsynaptic genes was also observed to be downregulated following *circHomer1* knockdown(22). It remains to be elucidated whether these mechanisms also contribute to the function of *circHomer1* in experience-dependent plasticity and the normal synaptic localization of Arc protein following MD.

The mechanism(s) of circRNA function are an area of active investigation. circRNAs are enriched in neurons and are believed to play critical roles in regulating neuronal development and synaptic plasticity(7–9, 33, 52). From within the top 100 candidates identified in our MD RNAseq screen, several circRNAs were already reported in the literature to have biological functions in other contexts(53–55). These include the neuronally-enriched circRNA *Cdr1as* (also known as ciRS-7), which has been found to contain a large number of binding sites for miR-7, a regulator of neurodevelopment(6, 56–59). This enrichment for miR-7 binding sites effectively serves as a “miRNA sponge”, and *Cdr1as* knockout mice exhibit behavioral deficits and altered neuronal electrophysiological properties(6, 8).

This study identified tens of circRNAs differentially expressed following experience-dependent plasticity and highlights *circHomer1*, a neuron-enriched circRNA derived from the Homer1 gene, as a critical regulator of V1 plasticity during the critical period. Our findings advance the understanding of circRNA regulation during experience-dependent plasticity and shed light on their functional significance in developmental processes.

## Acknowledgments

We thank Jamie Benoit for his contribution to the initial RNA-seq experiment. We thank Taylor Johns, Austin Sullins and other members of the Sur lab and the Ip lab for their help and support. We thank Hovy Wong for comments on the manuscript. We thank Lorena Rubino for her assistance in analyzing RNA-seq data. We thank Cara Kwong for her technical assistance in neuronal cultures. We thank the MIT BioMicroCenter for running the RNA-seq library preparation and sequencing. We thank scidraw.io for the schematic, licensed under a Creative Commons 4.0 license (https://creativecommons.org /licenses /by/4.0/). This work was supported by NIH grants R01MH085802 (M.S.), R01EY028219 (M.S.), F31EY033649 (K.T.), F32EY032756 (K.J.), the Simons Foundation Autism Research Initiative through the Simons Center for the Social Brain, MIT (M.S.), the National Research Foundation of Korea (RS-2023-00264980; T.K.), the Hong Kong Research Grants Council Early Career Scheme (24117220; J.I), General Research Fund (14117221; J.I.), Area of Excellence Scheme (AoE/M-604/16; J.I., A.F.), Theme-based Research Scheme (T13-605/18-W; J.I., A.F.), Lo Kwee-Seong Biomedical Research Fund (J.I.), Faculty Innovation Awards from the Faculty of Medicine CUHK (FIA2020/A/04; J.I.), and NARSAD Young Investigator Grant from the Brain & Behavior Research Foundation (J.I.).

## Author Contributions

J.I., N.M., M.N., M.S., K.J. conceived experiments with input from others. M.N., C.D. and S.Y. performed the cortical dissections and RNA and protein purifications. C.D., M.N and Y.Z. analyzed RNA-seq data. M.N., Y.C. and C.D. performed qPCR experiments. K.J., K.T., J.Z. and J.I. performed surgeries and viral injection, and carried out *in vivo* optical imaging experiments. K.L. analyzed optical imaging data. Y.C., S. Y. and J.Z. performed spine morphology analysis. Y.C., S.Y., T. K., D. Y. and J.I. performed eMAP experiments with inputs from K.C. J.I. performed the surface biotinylation assay and protein analyses. J.S. performed electrophysiological recordings and analyses. Y.C., Y.W., Y.D. performed neuronal culture experiments and relevant analyses. A.F. provided expertise on neuronal culture experiments. N.M. shared the sh-RNA constructs. G.H. performed the modeling of RNA expression. K.J., J.I., and M.S. contributed to analysis of experiments and interpretation of results. J.I., K.J. and M.S. wrote the manuscript with input from others. All authors edited the manuscript.

## Data, Materials, and Software Availability

All RNA sequencing data have been deposited in GEO. The code will be made available upon reasonable request.

## Figure Legends

Supplementary Figure 1. Transcriptomic analyses after MD in mouse V1.

(A) GO enrichment of the parental genes of differentially-expressed circRNAs after 3-day MD. DEG, differentially-expressed gene. (B) Volcano plot of mRNA sequencing of V1 after 7-day MD (n = 4 mice per group). (C) Volcano plot of circRNA sequencing of V1 after 7-day MD (n = 4 mice per group). (D) GO enrichment of the parental genes of differentially-expressed circRNAs after 7-day MD. (E) Fold change of differentially-expressed circRNAs and their linear isoforms after 7-day MD.

Supplementary Figure 2. sh-circHomer1 specifically targets *circHomer1* without affecting linear *Homer1* mRNA.

(A) sh-circHomer1 specifically reduces expression of *circHomer1* but not linear *Homer1* in mouse V1 (n = 4 mice per group, Student’s t-test). (B) sh-circHomer1 has no effect on total Homer1 protein levels as measured by Western blot (n = 3 mice per group). Data are presented as mean ± SEM. *p < 0.05.

Supplementary Figure 3. Distribution of spine volume on apical dendrites of layer 2/3 pyramidal neurons in V1 during MD.

Distribution of spine volume in (A) No MD, (B) 3-day MD, and (C) 7-day MD conditions (n = 235-334 dendritic spines from 3 mice per group, values were corrected assuming a 3^3^ expansion factor).

Supplementary Figure 4. Characterization of the effects of *circHomer1* depletion on membrane properties in layer 2/3 neurons of binocular V1.

(A) Representative traces show the response of sh-scramble or sh-circHomer1 infected neurons to −100, 0, and 300 pA current steps. The duration of each current step was 250 ms. (B) Quantification of the number of action potential spikes per current injection step shows the input-output relationship in sh-scramble and sh-circHomer1 infected neurons. (C) Rheobase, defined as the minimum amount of current injected to elicit action potential firing, was measured in response to the current injection protocol (n=7-10 neurons per group, Student’s t-test). (D-F) Passive membrane properties were measured in sh-scramble and sh-circHomer1 infected neurons (n=7-10 neurons per group, Student’s t-test). Data are presented as mean ± SEM.

Supplementary Figure 5. Characterization of the effects of *circHomer1* depletion on synaptic properties in layer 2/3 neurons of binocular V1.

(A) A representative image of the injection site shows an area of lentivirus-infected neurons in red, and in green three biocytin-filled cells that were recorded. The dotted line along the pial surface indicates the outline of the imaged slice. Scale bar = 100 µm. (B) Left, representative traces recorded from neurons in layer 2/3 of V1 expressing sh-scrambled (top) or sh-circHomer1 (bottom) lentivirus. Right, all events detected were scaled and averaged to show the average mEPSC kinetics. (C-D) Quantification of mEPSC amplitude (C) and frequency (D) in sh-scramble or sh-circHomer1 infected neurons (n=10 neurons per group, two-sample Kolmogorov-Smirnoff test). Data are presented as mean ± SEM. **p < 0.01.

Supplementary Figure 6. Alterations of surface AMPA receptor expression after *circHomer1* depletion.

(A) Expression of *circHomer1* (left), *c-fos* (middle), and *Homer1a* (right) RNAs following bicuculline treatment in cultured cortical neurons (n=4-5 per group, one-way ANOVA following Tukey multiple comparisons). (B) Representative images of cultured hippocampal neurons’ dendrites co-transfected with SEP-GluA1 (sGluA1) together with sh-scramble (top) or sh-circHomer1 (bottom) at DIV 11. Neurons were treated with bicuculline (20 µM) for 48 hours at DIV 17, and fixed at DIV 19. Con, control treatment; Bic, bicuculline treatment. Scale bar = 5 µm. (C) Percentage of sGluA1-positive spines under different conditions. For each neuron, 3 segments of dendrite with length of 30 µm were randomly selected, and the average percentage were calculated (n=12 neurons per group, two-way ANOVA following Tukey multiple comparisons). *: compared to sh-scramble: Con; ^†^: compared to compared to sh-circHomer1: Con. Data are presented as mean ± SEM. *p < 0.05; **, ^††^p < 0.01; ***p < 0.001.

Supplementary Figure 7. Modeling of *circHomer1* expression.

(A) Schematic of the model components for each experiment. Arrows represent rates fit in each model of either transcription, circularization, or degradation. These values were fit for each model and experiment combination separately to minimize squared error from measured *Homer1a* and *circHomer1* levels. (B) Simulated levels of *circHomer1* RNA for the constant circularization (pink), variable circularization (red), and full models (dark red) for monocular deprivation (left), development (middle), and bicuculline (right) experiments. The measured, normalized levels of *circHomer1* in each experiment are overlaid (black dots). (C) Normalized Akaike information criterion (AIC) scores for each model tested. (D) Expanded results from the full models, showing the simulated levels of both *circHomer1* and *Homer1a* RNA (black and purple lines, respectively) overlaid with the experimentally measured levels of *circHomer1* and *Homer1a* RNA (black and purple dots, respectively) for each experiment.

Supplementary Table 1: RNAseq results of mRNAs detected from 3-day MD experiment.

Supplementary Table 2: RNAseq results of circRNAs detected from 3-day MD experiment.

Supplementary Table 3: RNAseq results of mRNAs detected from 7-day MD experiment.

Supplementary Table 4: RNAseq results of circRNAs detected from 7-day MD experiment.

Supplementary Table 5: Primers list.

## Materials and Methods

### Animals

Experiments were carried out in mice under protocols conforming to NIH guidelines and approved by MIT’s Animal Care and Use Committee or by CUHK’s Animal Experimentation Ethics Committee. Wildtype C57BL/6J mice (JAX000664) were sourced from Jackson Laboratory or CUHK’s Laboratory Animal Services Centre. Mice were group-housed whenever possible with up to 5 same-sex mice per cage. The cages were in a standard animal facility room with a 12-hour/12-hour light/dark cycle. Food and water were available ad libitum. Both male and female mice were used for experiments.

### Eyelid suture and monocular deprivation

For monocular deprivation experiments, mice were anesthetized using 2-3% isoflurane. The top and bottom eyelid of the right eye was trimmed, and sterile nylon sutures (7–0) were used to suture the eyelid, which were further sealed with Vetbond (3M). The suture was inspected for the next 3-7 days post-closure to ensure that the eye did not reopen. Mice that exhibited incomplete eyelid closure were removed from the experiment.

### *circHomer1* depletion via shRNA and validation

For *circHomer1* depletion experiments, the following viruses were used: lentivirus expressing pLV-mU6-scrambled-shRNA::SYN-tdTomato or pLV-mU6-*circHomer1*a-shRNA::SYN-tdTomato (1.2×10^9^ IFU/mL, System Biosciences), or AAV9 expressing pAV-U6-sh-scramble-CMV-DsRed or pAV-U6-sh-circHomer1-CMV-DsRed (2×10^13^ vg/mL, Vigene). For the *in vivo* shRNA experiments, binocular V1 in the left hemisphere of P15 pups was targeted. Briefly, sh-scrambled or sh-circHomer1 viruses were loaded into a pulled-glass micropipette with a beveled tip, lowered into L2/3, and virus was infused at a rate of 100 nL/min. The following coordinates were used (in mm from lambda at P15): AP: 0.5 to 1.0; ML: −2.8, DV :-0.2 to −0.25. A minimum of 10 days was allowed for viral transduction and sufficient expression of the constructs in all experiments. To estimate the efficiency and specificity of sh-circHomer1 after virus injection, the mouse brain was examined using a dual fluorescent protein flashlight (Nightsea) after mouse sacrifice and dissection. the tissue expressing RFP was dissected for RNA purification for RT-qPCR (see below), and protein isolation for Western blot. For protein isolation, the RFP-expressing tissue were homogenized in RIPA buffer (1% NP40, 0.1% SDS and 0.5% sodium deoxycholate in PBS). The homogenate was centrifuged at 14,000 x g for 15 min and the supernatant was collected. Protein concentration was measured by BCA protein assay, and 20 µg total protein lysate was boiled for 5 min with 6x loading buffer. Standard Western blot was performed. Antibodies used include Homer1 (160003; Synaptic Systems), Homer1a (160013; Synaptic Systems), and α-Tubulin (sc32293; Santa Cruz).

### RNA Sample Preparation

RNA was extracted from fresh tissue. Tissue for RNA extraction was isolated from V1 contralateral and ipsilateral to the sutured eye after 3-day MD (at P28), 7-day MD (at P32) or from mice that had not undergone MD (various ages based on experiment). Briefly, mice were anesthetized with isoflurane and transcardially perfused with ice-cold phosphate buffered saline (PBS) for at least 5 min. A sterile scalpel blade and forceps were used to surgically micro-dissect the appropriate tissue section. The dissected tissue was immediately placed into a 2-mL lysing tube containing 1.4 mm spherical ceramic beads (MPBio, Lysing Matrix D, Cat. No. 116913050) and pre-filled with 1 mL of ice-cold TRIzol (Ambion, Cat. No. 15596-026) and then immediately homogenized with a bead-based homogenizer (FastPrep-24 5G, MPBio) using the preset settings for mouse brain tissue (8.0 m/sec, 30 sec). Once foaming has subsided at room temperature, the homogenate was transferred into 2 mL Phase Lock Tube (Heavy; 5Prime Bio, Cat. No. 2302830) pre-filled with 200 µL chloroform and briefly vortexed. Samples were centrifuged at 12,000 x g for 10 min at 2-4°C. The colorless upper aqueous layer was then transferred into a new 2.0 mL LoBind tube that was pre-filled with 1.5 volumes of 100% EtOH (∼600 µL), then mixed thoroughly by pipetting up and down several times, and then briefly centrifuged. The sample was furthered purified using Zymo RNA Clean & Concentrator-5 (Zymo Research, R1016) according the manufacturer’s recommended protocol and always performed with the optional in-column DNase I treatment. RNA was eluted twice with ≥ μ 1x TE buffer (pH 7.5). The approximate concentration and relative purity of RNA was determined using a NanoDrop 2000 spectrophotometer (Thermo Scientific). The purified RNA was either immediately used for downstream applications (i.e., reverse transcription) or stored at −80°C.

### RNA sequencing library preparation

For samples to be sequenced, RNA quality was assessed using AATI Fragment Analyzer. Samples with RNA Quality Number of >8.0 were further processed. Indexed cDNA libraries were generated using the SMARTer Stranded Total RNA-Seq Kit v2 and multiplexed sequencing was performed on Illumina HiSeq 2000 or Novaseq 6000.

### RNAseq Analysis

For the mRNA sequencing, read alignment was performed using HISAT2(60). Fragment counts were obtained using featureCounts(61). Differential expression analysis was performed using DESeq2(62). For the circRNA sequencing, reads were aligned and fragment counts were obtained using circtools(23). Fragment counts were obtained using the Cufflinks pipeline(63). Differential expression analysis was performed using the Bioconductor package Limma(64). Gene Ontology enrichment analysis was performed using the Bioconductor package clusterProfiler(65). Differential expression analysis of 3-day MD RNAseq data was performed comparing the contralateral side and ipsilateral side within animal. For 7-day MD RNAseq, differential expression analysis was performed comparing control, no MD animals and MD animals (contralateral side).

### cDNA Preparation for qPCR

Total RNA was reversed transcribed using SuperScript IV VILO with ezDNase treatment (Thermo Fisher, Cat. No. 11766050) according to manufacturer’s recommended protocol with the following modifications: the maximum input of total RNA was lowered to 2 µg per 20 µL reaction, ezDNase digestion was extended to 5 min, and reverse transcription temperature was increased to 55°C. After the RT reaction, cDNA was diluted with 1x TE buffer (pH 8.0).

### qPCR Analysis

Applied Biosystems QuantStudio 3 Fast thermocycler was used for all qPCR amplification and detection in this study. PowerUp SYBR Green Master Mix (Applied Biosystems, Cat. No. A25742) was used according to the manufacturer’s recommended protocol. Primers used were listed in Supplementary Table 5. Mouse *Gapdh* was used as the internal control for normalization. For the fold change calculation, the ΔΔ method was used.

### Optical imaging of intrinsic signals

At postnatal day P15, sh-scramble or sh-circHomer lenti-viral vectors were injected into left binocular V1 as described above, and at P22 a 3 mm craniotomy was performed over the same area, where a 5 mm diameter glass window stacked on top of a 3 mm diameter glass window was fitted over the craniotomy with dental cement, together with a metal headplate. Baseline optical imaging was performed at P25, prior to MD. MD lasted either 3 days or 7 days, after which sutures were removed and the closed eye was reopened under isoflurane anesthesia before optical imaging experiments to assess the effects of 3-day or 7-day MD on V1 responses. Optical imaging was performed as previously described(26). Briefly, mice were lightly anesthetized with isoflurane (0.5-1%), the window was cleaned with 70% ethanol and a cotton tipped applicator, and the headplate was attached to the imaging rig to minimize head movements. Green light (560 nm) was used to focus 400 μ below the cortical surface. Red light (630 nm) was used for functional imaging, and the change in reflectance was captured by an electron multiplying CCD camera (Cascade 512B; Roper Scientific) during the presentation of visual stimuli.

### Visual stimulation and optical imaging analysis

The visual stimulus was a horizontal bar 30° wide, made of flickering checkerboard over a black background, drifting continuously through the peripheral–central dimension of the visual field. After moving to the last position, the bar would jump back to the initial position and start another cycle of movement; therefore, the chosen region of visual space (72° x 72°) was stimulated in a periodic manner (12 s/cycle, 20 repetitions). Images were continuously captured at the rate of 30 frames/s during each stimulus session of 4 min, with a separate stimulus session for each of the 4 cardinal directions. A temporal component at the stimulus frequency (12s^-1^) was calculated pixel by pixel from the whole set of images using custom python scripts (https://github.com/Palpatineli/oi_analyzer). The amplitude of the FFT component was used to measure the strength of visually driven response for each eye, and the ODI was derived from the response (R) of each eye at each pixel as ODI = (R_contralateral_ - R_ipsilateral_)/(R_contralateral_ + R_ipsilateral_). The binocular zone was defined as the cortical region that was driven by both eyes. The response amplitude for each eye was defined as fractional changes in reflectance over baseline reflectance (ΔR/R × 10^-3^), and the top 50% pixels were analyzed to avoid background contamination.

### Epitope-preserving magnified analysis of the proteome (eMAP)

Mice were injected at P15as described above with AAV2/9-hSyn-DIO-EGFP (1×10^13^ vg/mL), AAV2/9-CaMKIIa-Cre (5×10^8^ vg/mL), and AAV9 expressing either pAV-U6-sh-scramble-CMV-DsRed or pAV-U6-sh-circHomer1-CMV-DsRed (1×10^13^ vg/mL). Animals were allowed to express virus for a minimum of 10 days before MD. After MD, mice were transcardially perfused under deep anesthesia with ice-cold PBS followed by ice-cold 4% PFA (in PBS), and then brains were post-fixed in 4% PFA at 4°C overnight. eMAP processing was performed following protocols previously described(66, 67). Tissue embedding was performed using eMAP solution (30% acrylamide, 10% sodium acrylate, 0.1% bisacrylamide, 0.03% VA-044 in PBS) under vacuum, following hydration using hydration solution (0.02% sodium azide in PBS) and sectioning into 60um thick using Leica VT1000S vibratome. Tissue clearing was performed by incubation in clearing solution (6% SDS, 0.1 M phosphate buffer, 50 mM sodium sulfite, 0.02% sodium azide in DI water, pH 7.4) at 37°C for 4 hours. If applicable, after washing with PBST (0.1% Triton X-100, 0.02% sodium azide in PBS), samples were stained with primary antibodies against GFP (A10262, Invitrogen) and Arc (156003, Synaptic Systems) diluted in PBST at 37°C for 48 hours. After washing with PBST, samples were stained with secondary antibodies diluted in PBST for 48 hours. Expansion was performed using 0.01× PBS before imaging. Approximately 3× total linear expansion was achieved consistently. Pyramidal neurons in V1 which were double-positive for RFP and GFP were identified and imaged using Leica SP8 confocal microscope with 20× 0.75 NA or 63× 1.2 NA water immersion lens.

### Spine morphology and Arc puncta volume analysis

Dendrites and dendritic spines for analysis were selected and analyzed by an experimenter blinded to experiment condition. To analyze dendritic spine morphology, the length (L), head width (H) and neck width (N) were measured by ImageJ. Dendritic spines were classified into 4 types according to the criteria previously described(29). Briefly, the average length of spine head in the No MD sh-scramble group (H̅) was calculated. Protrusions with H>N and H> H̅ were defined as mushroom spines; protrusions with H>N and H<H̅ were defined as thin spines; protrusions with H≤N and L< H̅ were defined as stubby spines, and protrusions with H≤N and L>1.5* H̅ were defined as filopodia. For dendritic spine volume and surface area analysis, 3D reconstruction and quantification of dendritic spines volume was performed using the “Filament” function in Imaris software. The analysis of Arc puncta volume was performed using the “Surface” function in Imaris, and only the Arc puncta located at dendritic spines were selected for quantification. The raw data was then divided by 3^3^ (=27) to convert to values that closely approximates the unexpanded volume.

### Preparation of acute brain slices for electrophysiology

Mice injected with either sh-scramble or sh-circHomer1 lentivirus (as described above) were anesthetized with 2-3% isoflurane and decapitated. Brains were rapidly removed and placed into ice-cold low-Ca2+, low-Na+ sucrose cutting solution consisting of (in mM): 234 sucrose, 11 glucose, 24 NaHCO3, 2.5 KCl, 1.25 NaH2PO4, 10 MgSO4, and 0.5 CaCl2. Cutting solution was oxygenated with a mixture of 95% O2 and 5% CO2. 300µM slices were cut in a coronal orientation using a vibratome (Leica Biosystems, Buffalo Grove, IL) and placed into a recovery chamber filled with oxygenated ACSF consisting of (in mM): 126 NaCl, 2.5 KCl, 1.25 NaH2PO4, 1 MgSO4, 2 CaCl2, 10 glucose, 26 NaHCO3. Slices were allowed to recover at 34°C for 1 hour before being kept at room temperature (25°C) for the remainder of the experiment.

### Current clamp recordings and analysis

TdTomato-positive neurons were identified in Layer 2/3 of the V1 binocular region. Recordings were made at 34°C in ACSF containing 10 µM CPP, 20 µM DNQX, and 10 µM SR95531 to isolate passive membrane properties. Signals were collected at 10kHz using a Multiclamp 700B amplifier and digitized using a Digidata 1440A (Molecular Devices, San Jose, CA). Whole-cell patch clamp recordings were made using borosilicate glass electrode with an open-tip resistance of 2-5 MΩ filled with an intracellular solution consisting of (in mM): 130 potassium gluconate, 10 HEPES, 5 KCl, 5 EGTA, 2 NaCl, 1 MgCl2, 10 TEA-Cl, 10 phosphocreatine, 2 Mg-ATP, 0.3 Na-GTP (pH adjusted to 7.28, osmolarity adjusted to 290 mOsm). Current steps of 250 ms duration were applied, from −100 pA increasing to 650 pA at 25 pA increments. The holding current was returned to 0 pA for 750 ms between each current injection step. Ra (access resistance) and Rm (membrane resistance) were also measured in response to a 5 mV depolarizing step, and membrane capacitance was calculated as 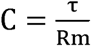, where τ is the decay of the capacitive transient in response to the 5 mV step. Access resistance was monitored throughout the recording and recordings with >20% change in access resistance or had access resistance >30 MΩ were excluded.

### mEPSC recordings and analysis

Slices were prepared as above. Recordings of tdTomato positive neurons were made at 34°C in ACSF containing 10 µM CPP, 10 µM SR95531, and 1 µM tetrodotoxin to isolate miniature AMPA receptor-mediated currents. Signals were collected at 10kHz and filtered at 2kHz using a Multiclamp 700B amplifier and digitized using a Digidata 1440A (Molecular Devices, San Jose, CA). Whole-cell patch clamp recordings were established using borosilicate glass electrode with an open-tip resistance of 2-5 MΩ filled with an intracellular solution consisting of (in mM): 100 potassium gluconate, 10 HEPES, 20 KCl, 0.5 EGTA, 10 NaCl, 8 phosphocreatine, 2 Mg-ATP, 0.3 Na-GTP (pH adjusted to 7.23, osmolarity adjusted to 290 mOsm). Neurons were voltage-clamped at −70 mV and stable gap free recordings were made for at least 2 minutes. Access resistance was monitored throughout the recording and recordings with >20% change in access resistance or had access resistance >30 MΩ were excluded. mEPSC events were detected using the MiniAnalysis software (Synaptosoft, Fort Lee, NJ).

### Surface Protein Biotinylation Assay

For the measurement of surface proteins, we first prepared 300 µm acute coronal slices containing V1 in ice-cold ACSF and then washed slices 3 times in ice-cold ACSF in 6-well plate on shaker. The sections were incubated in 100 µM S-NHS-SS-biotin for 45 min (∼1 mL per well). The superficial layers of V1 were dissected into a new 1.5 mL tube filled with ice-cold ACSF and homogenize in RIPA buffer. The homogenate was centrifuged at 14,000 x g for 5 min and the supernatant was transferred into a new 1.5 mL tube. ∼20% volume was transferred into a new 1.5 mL tube and stored at −80°C to be used later as a total protein control. ACSF was added to the remaining supernatant for a final volume of 1 mL per sample. 40 µL of streptavidin beads were added and incubated on a shaker overnight at 4°C. Samples were then centrifuged at 3,500 x g for 1 min. The supernatant was discarded and then the beads were washed 3x in 1:1 v/v ice-cold solution of ACSF and RIPA buffer. Then, 40 µL of 2x loading buffer was added, mixed briefly, and boiled for 5 min. Standard Western blot was performed. Antibodies used include GluA1 (04-855, clone C3T; Millipore), and pan-Cadherin (ab6529; Abcam).

### Neuronal cultures, transfection, and pharmacological treatment

Cultured primary rat hippocampal and cortical cells were prepared as previously described(68). In brief, Sprague–Dawley rat embryos were sacrificed on embryonic day 18. The hippocampus and cortices were dissected, and the cells were dissociated with trypsin. Hippocampal cells were cultured on 18-mm coverslips coated with 1 mg/mL poly-D-lysine at 1 × 10^5^ cells per coverslip in Neurobasal Plus medium supplemented with 2% B27 Plus and 0.5mM L-glutamate. Cortical cells were cultured on 60-mm cultural dishes coated with 100μg/mL poly-L-lysine at 3 × 10^6^ per dish in Neurobasal medium supplemented with 2% B27 and 10mM glucose. All cells were maintained at 37 °C in a humidified atmosphere with 5% CO_2_. The cortical neurons were treated with bicuculine (20 μM) at DIV 13. Total RNA of cortical neurons was extracted at 0, 2, 6, 24, or 48 hours later using NucleoSpin RNA Mini kit (Macherey-Nagel, 740955) according to the manufacturer’s protocol. The hippocampal neurons at DIV 11 were transfected with a *circHomer1* or scrambled control shRNA construct and SEP-GluA1 using calcium phosphate precipitation(68). The transfected neurons were then treated with bicuculine (20 μM, 48 h) for subsequent morphological analysis. At DIV 19 the hippocampal neurons were fixed with 4% PFA, and blocking was performed for 1 hour at room temperature with 1% BSA in PBS. Antibody against GFP (A10262, Invitrogen) was diluted in 1% BSA in PBS and incubated with cells at 4°C overnight. After washing with PBS, the cells were incubated with secondary antibody for 1 hour at room temperature. The cells were then washed with PBS and mounting was performed. Imaging was performed under Leica SP8 confocal microscope using a 63x oil immersion lens with 1.4 numerical aperture.

### Computational modeling

As it is unclear what stages in the biogenesis of *circHomer1* confer its activity-dependence, we compared three nested dynamical system models(69) of *circHomer1* expression with the same basic architecture to assess which best fit our data. We modeled three time-varying outputs: (1) *Homer1b* mRNA levels (H1b), (2) *Homer1a* mRNA levels (H1a), and (3) *circHomer1* RNA levels (CH). Dynamical changes in gene expression were described by the following differential equations:

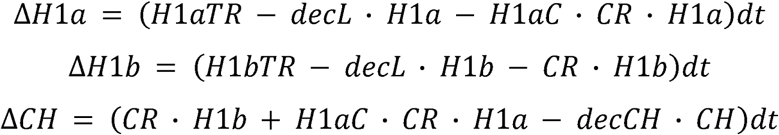

with H1a/bTR denoting transcription rates of pre-mRNA, decL denoting linear mRNA decay rate, decCH denoting circular RNA decay rate, CR denoting circularization rate, and H1aC denoting a logical operator of whether *Homer1a* can be circularized (1 if true, 0 if false). Parameters were fit separately for each model/experiment combination, but the fit parameter values for a particular model were largely consistent across experiments.

We hypothesized(21, 47) that the activity-dependence of *circHomer1* expression was most likely to arise from circularization of an activity-dependent pre-mRNA (*Homer1a*) and/or activity dependent circularization of a constitutively expressed pre-mRNA (*Homer1b*). The three models tested were as follows. (1) The constant circularization model fit a constant circularization rate of pre-*Homer1a* and pre-*Homer1b*, with the only time varying parameter being the transcription rate of *Homer1a* which was fit to measured data. (2) The variable circularization model allowed the circularization rate to vary over time, however only pre-*Homer1b* could be circularized. (3) The third, full model also included time varying circularization but both pre-*Homer1b* and pre-*Homer1a* could be circularized. *Homer1a* and *circHomer1* levels were explicitly fit to minimize squared error from observed datapoints for each experiment. *Homer1b* levels were treated as a ‘hidden’ variable; as they were not measured experimentally and were not a part of model testing.

Models were initialized 10 days before the first timepoint to allow RNA levels to stabilize before permitting any changes in transcription or circularization. Because of the varied measurements and methods across experiments, data in each model were normalized to the first timepoint, T=0. In models where circularization of pre-*Homer1a* was permitted, the same circularization rate was applied to both pre-*Homer1a* and *Homer1b* pools. Fit circularization rates were 1 to 2 orders of magnitude smaller than the linear splicing rates(70), and varied with activity by at most a factor of 4. In each of the models, the decay rate of *circHomer1* was constrained to be slower than that of linear *Homer1* mRNA by a factor of 2 ≥ (48). All models were run with 15 minute timesteps. *Homer1b* transcription rate, linear decay rate, and circular decay rate were fit as constant parameters in each model. *Homer1a* transcription rate was fit as a variable parameter in each model. Circularization rate was fit as a constant parameter in the constant circularization model, and a variable parameter in the variable circularization and full model. Fitting was performed and AIC scores for each model were computed using the python package lmfit.

### Statistical analysis

Data are presented as mean ± SEM in quantitative analysis unless other specified. When two independent experimental groups were analyzed, Student’s t-test was performed, while paired t-test was performed when two paired experimental data were analyzed. When more than two independent experimental groups were analyzed, one-way ANOVA followed by Tukey post hoc test was performed. For those experiments exploring the effect of *circHomer1* during MD by *circHomer1* depletion, two-way ANOVA followed by Tukey post hoc test was performed. For optical imaging, mixed-effects model following Holm-Sidak multiple comparisons test was performed. For mEPSC quantification, two-sample Kolmogorov-Smirnoff test was performed. Statistical significance was defined as p < 0.05.

