## Supplementary figures and images for "The noncoding circular RNA *circHomer1* regulates synaptic development and experience-dependent plasticity in mouse visual cortex"

### Supplementary Figure 1

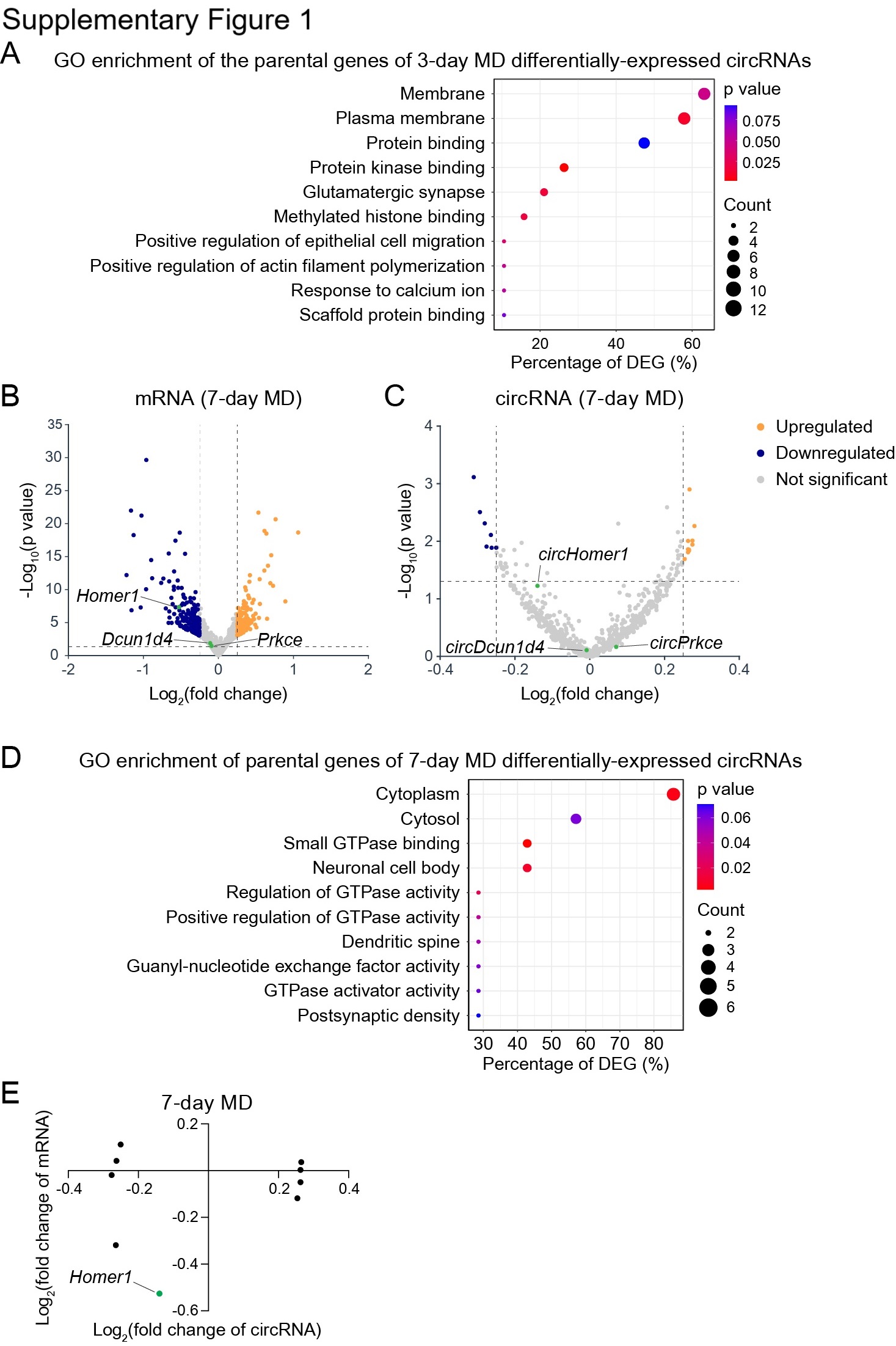

### Supplementary Figure 2

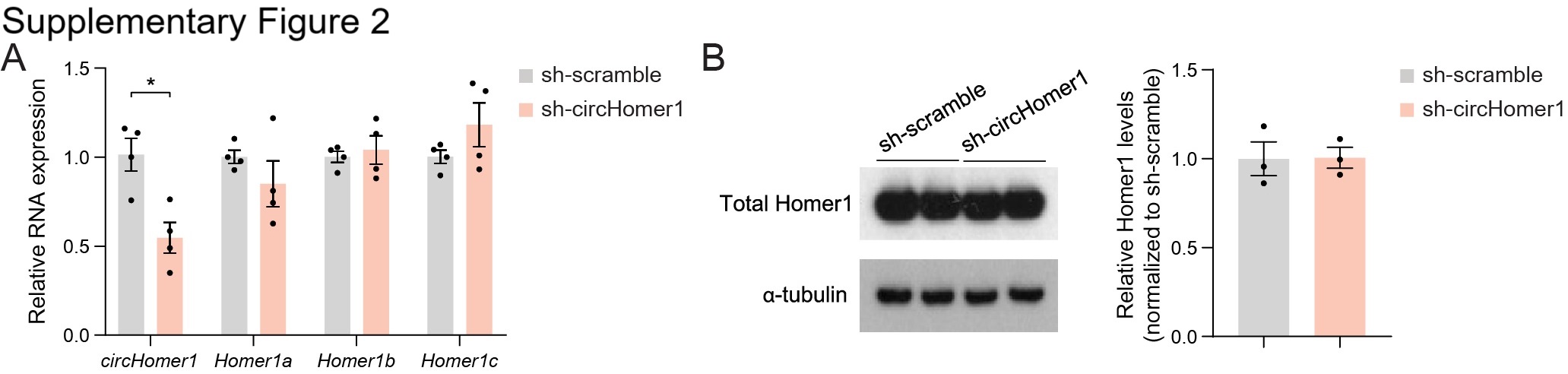

### Supplementary Figure 3

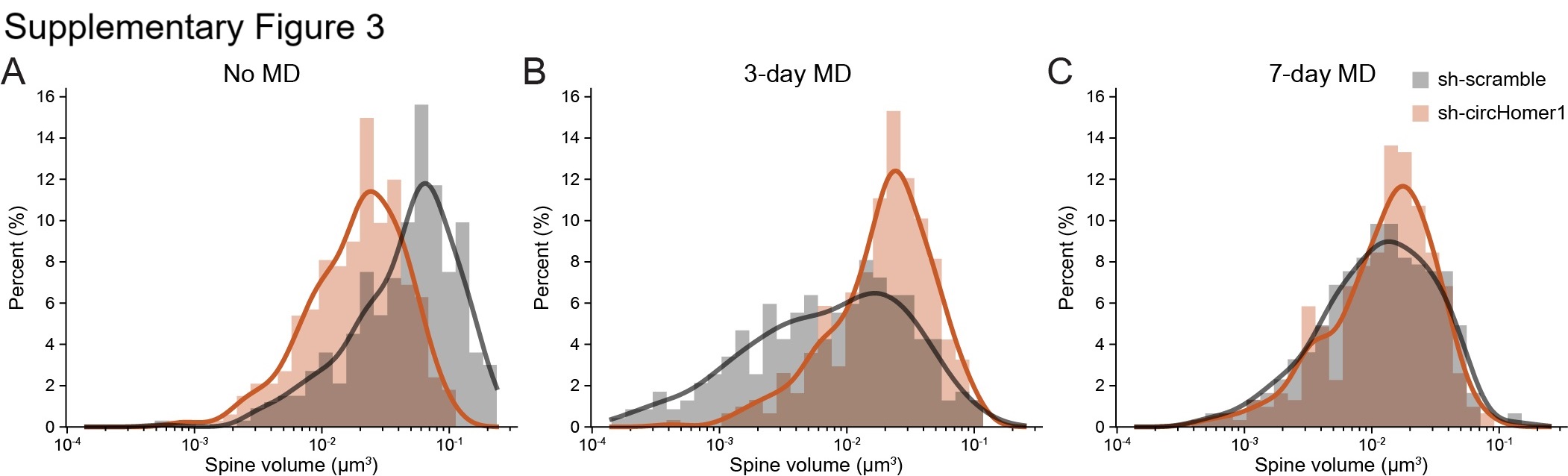

### Supplementary Figure 4

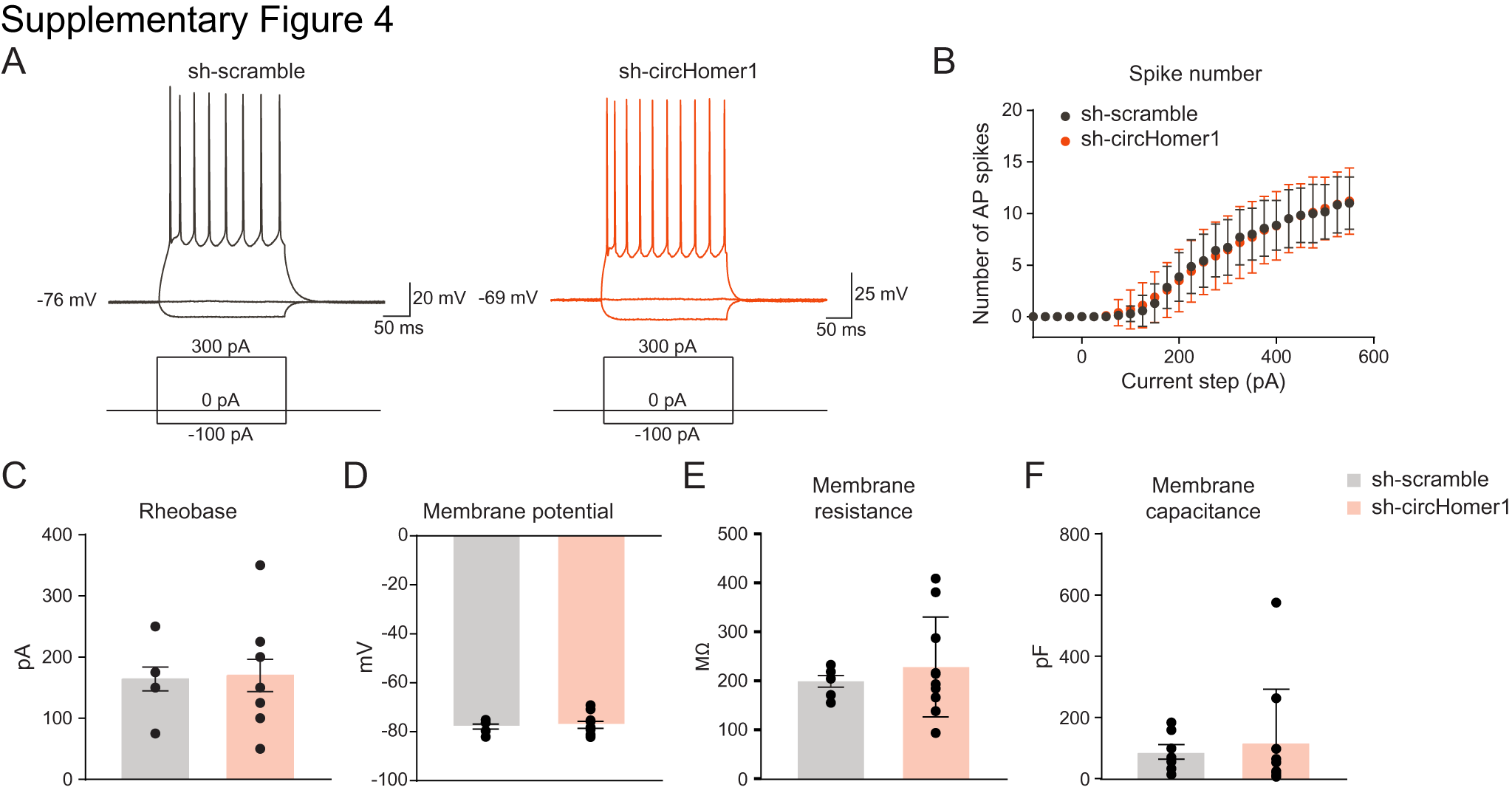

### Supplementary Figure 5

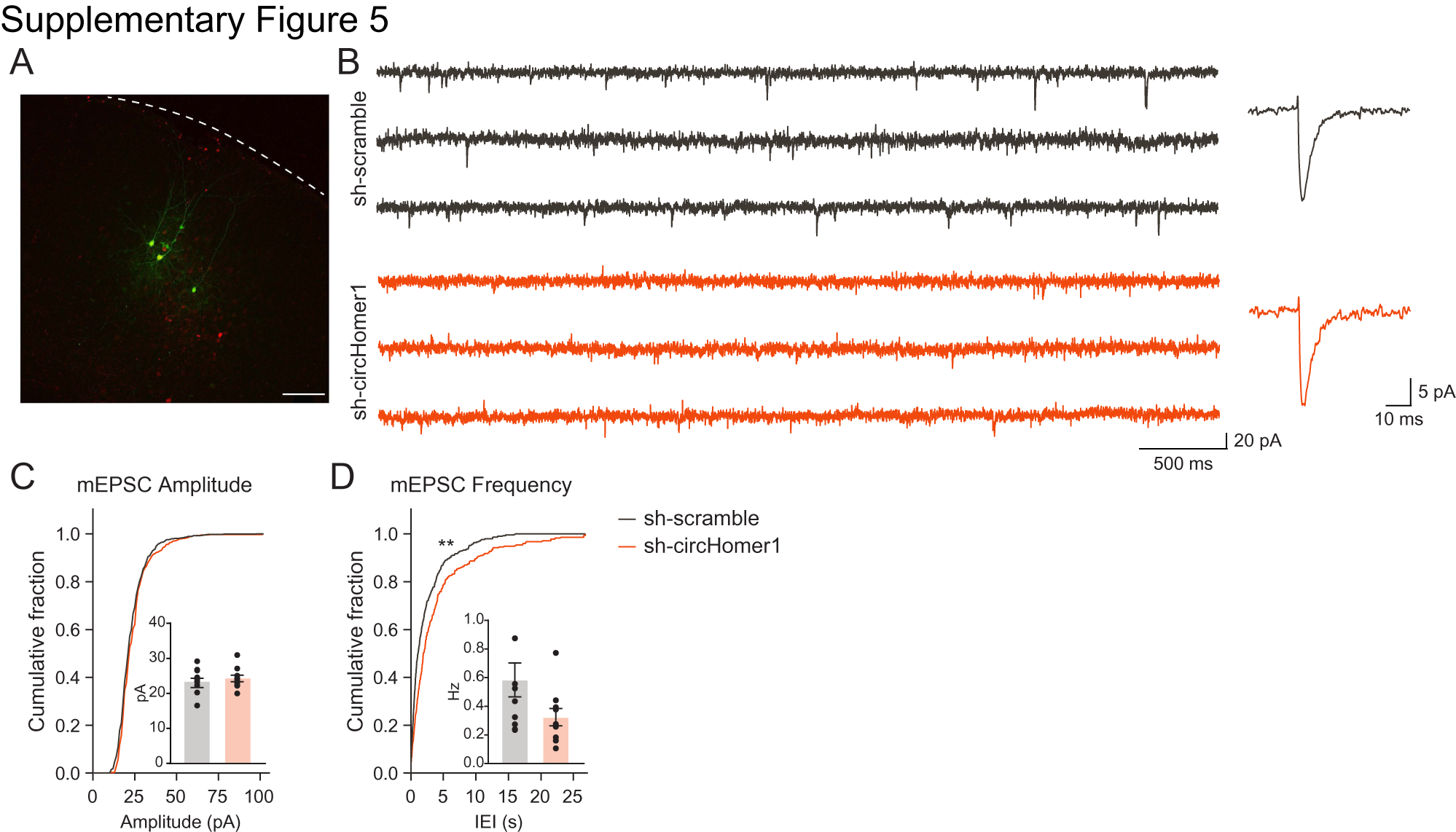

### Supplementary Figure 6

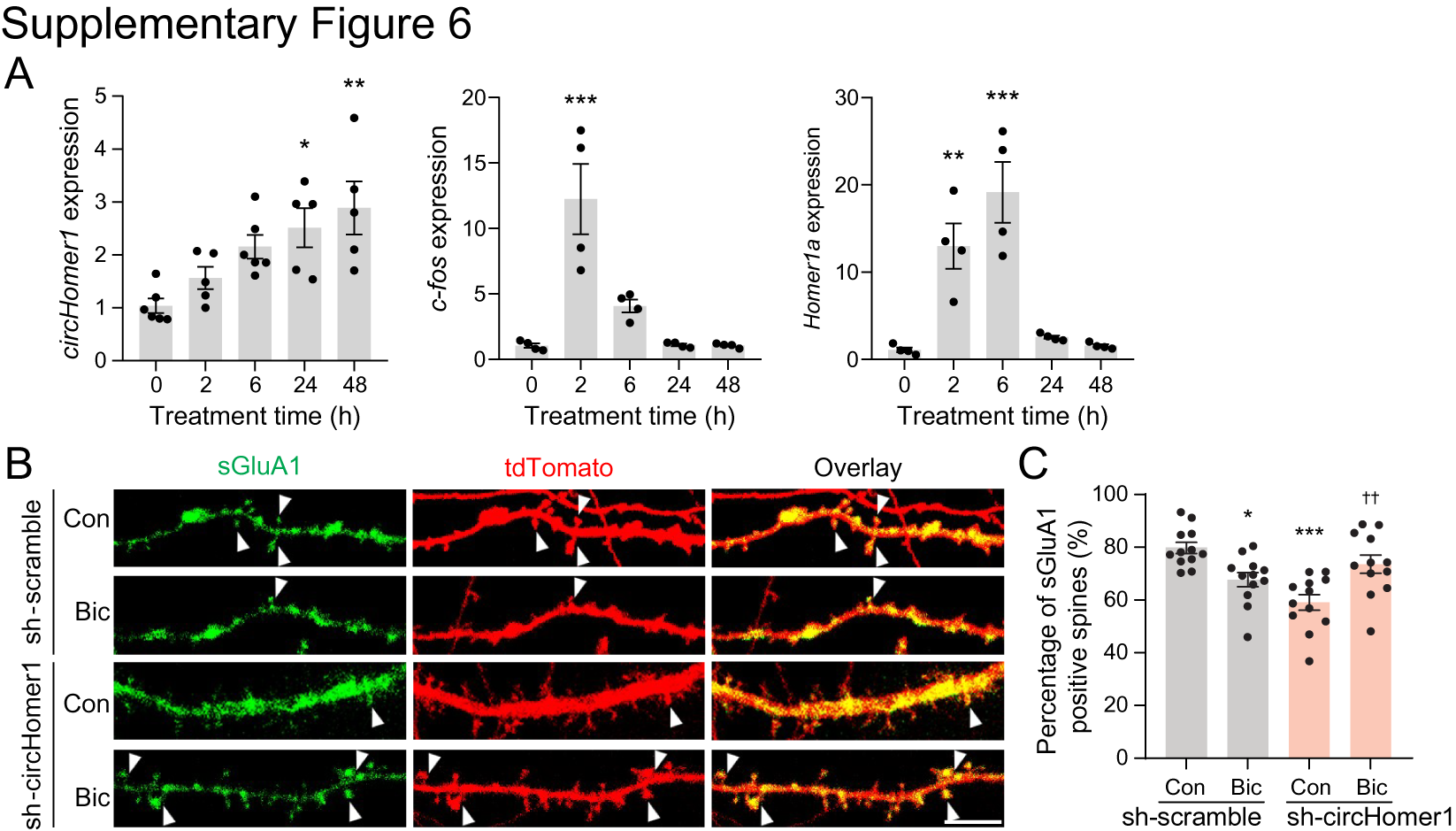

### Supplementary Figure 7

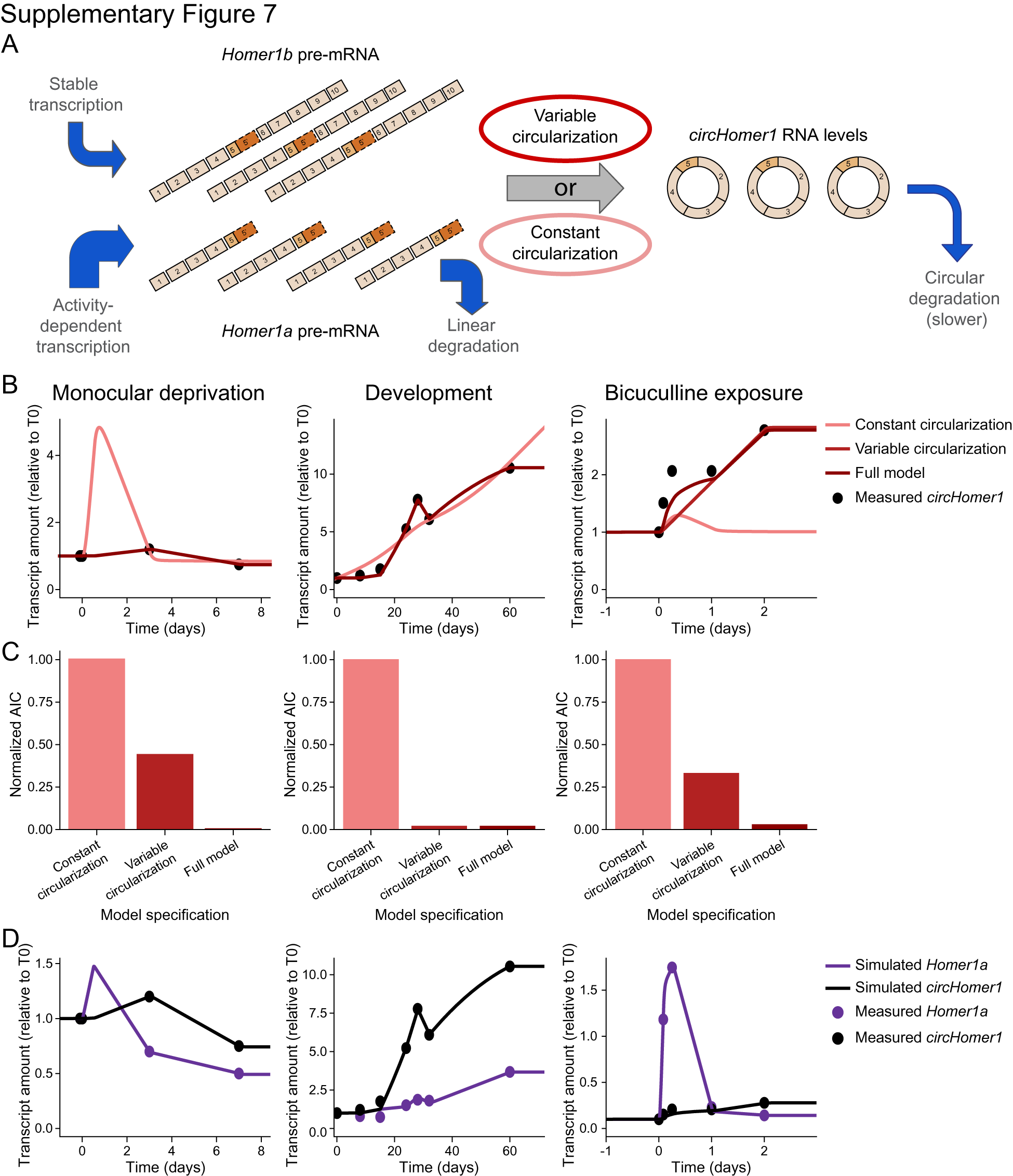
